# Microbial sedimentary DNA from a cultural landscape disentangles the impacts of humans and nature over the past 13.5 thousand years

**DOI:** 10.1101/2025.03.14.642998

**Authors:** Yi Wang, David Schleheck, Elena Marinova, Martin Wessels, Sebastian Schaller, Flavio S. Anselmetti, Antje Schwalb, Mikkel W. Pedersen, Laura S. Epp

**Affiliations:** Limnological Institute, Department of Biology, University of Konstanz, Konstanz, Germany; Laboratory of Archaeobotany, State Office for Cultural Heritage Baden-Württemberg, Gaienhofen-Hemmenhofen, Germany; LUBW State Institute for Environment Baden-Württemberg, Inst. for Lake Research, Langenargen, Germany; Institute of Geological Sciences and Oeschger Centre for Climate Change Research, University of Bern, Bern, Switzerland; Institute of Geosystems and Bioindication, Technische Universität Braunschweig, Braunschweig, Germany; Centre for Ancient Environmental Genomics, Globe Institute, University of Copenhagen, Copenhagen, Denmark

**Keywords:** Paleoecology, sedimentary DNA, environmental DNA, bacteria, archaea, ancient prokaryote DNA, Lake Constance, network analysis, DNA damage

## Abstract

Bacteria and archaea are currently under-characterised in palaeoecological studies, despite their ubiquity, high diversity and tight integration with the biotic and abiotic environment and human activity. The complexity of their assemblages, and the difficulties in separating living- from paleo-prokaryotes renders analyses challenging. Here we present an ancient prokaryote metagenomic time-series from a sediment core of Lake Constance, a large and deep perialpine lake from temperate Europe, spanning the last 13,500 years of natural and anthropogenic impact. We mapped DNA to reference genomes and estimated the DNA damage of taxa, which displayed a monotonic relationship with time. By constructing co-abundance networks we recognize major microbial assemblages, containing both ancient and living microbes, that show specific dynamics. Short-term and often low-abundance assemblages are linked to the Pleistocene-Holocene transition, floods and human activities. Noticeably, certain lineages harbouring microbes common in human-impacted environments expanded during the Middle Ages and Modern time. Some abundant taxa that were linked to various freshwater and soil environments persisted through millennia. By extricating various sources and trajectories of change, we demonstrate the power of prokaryotic sedimentary DNA in revealing long-term eco-evolutionary outcomes caused by both nature- and humans.

## Introduction

Prokaryotes exhibit remarkable diversity and metabolic versatility, and are tightly integrated with their environment. Found in soil, water, subsurface and eukaryotic hosts, they are copious in biomass and indispensable in the functioning of ecosystems they inhabit^1–4^. The crucial role of prokaryotes in shaping Earth’s primeval environment has been indicated by microbial fossils, such as organic-rich shales and stromatolites formed by Archaean age photosynthesizing bacteria^5^. For more recent environmental history and ongoing biogeochemical processes, the extraction of molecular remains such as DNA and lipids have become the major method to recover microbial diversity and function. Many studies have so far characterised active microbes from modern and ancient habitats of known functional significance, such as carbon reservoirs^6–8^, extreme environments^9^ and habitats under change^10,11^. A few studies have investigated sediments aged thousands and millions of years, with a focus on environmental history^12,13^. However, despite their ecological significance and close connections to the environment, paleoecological investigations of prokaryotes in the environment lag behind their eukaryotic, macroscopic counterparts.

Early studies extracted ancient bacterial DNA from sediments with success^14–16^, but the reliance on PCR-based methods, which do not provide independent measures for DNA degradation, means that the data could not differentiate ancient bacterial DNA from that of living microbes, be they endogenous to the sample or contaminants. This challenge of ancientness authentication^17,18^ continued to prevail in paleoecological studies that targeted prokaryotes with 16S rDNA amplicon sequencing^19–22^, a method that is widely used to recover microbial diversity^23^ but obliterates signals of age-dependent DNA degradation. One way to circumvent this issue is to focus on environmental or ecological processes that only involve specific lineages, which are, in the best cases, not sediment-dwelling^24,25^. Yet studies on prokaryotic DNA from ancient substrates face another issue that is of little concern for those that extract DNA from human remains^26–28^, that is, the sources of the DNA. Some studies interpreting microbial DNA in its entirety, suggested that prokaryotic communities in ancient substrates, living and perished, stem from complex sources ^29^, i.e. they may be correlated with paleoenvironments of local, depositional nature^20,30^, under regional, historical influence^31^ or show responses to climate changes at global scale^13,32^. However, even among studies that successfully quantified DNA degradation, very few have teased apart these different fractions of assemblages at a natural site^13,31,32^.

The explicit acknowledgement that microbial assemblages retrieved from paleoecological archives stem from communities of different origins but are then reassembled through depositional processes is currently lacking in prokaryote paleoecology. These communities consist of phylogenetically diverse members; among them, each might display independent dynamics over time. In modern microbial ecology, the concept of community has been explored with data-driven methods such as network analysis^33,34^, revealing for Earth’s microbiome not only patterns that connect taxa to the environment, but also niches, keystone species and community dynamics that are not driven by the environment^35,36^. This network approach is, however, rarely applied in the study of temporal dynamics of microbes^19,37^ even in modern datasets^37–40^. In paleoecology, it is further argued that co-occurrence networks more likely reveal death-assemblages^41^ formed by deposition pathways, rather than representing interacting communities. But beyond being an agent of bias, taphonomy is also ecological by nature^42–44^. This implies that a network perspective on the temporal distribution of microbial molecular remains may shed light on unknown paleoecological processes.

In this study, we address the aforementioned knowledge gaps by describing the diversity, origins and occurrence timelines of prokaryotes in a lake sediment succession spanning the last 13,500 years. We focus on taxonomic characterisation as ecological functions of prokaryotes are shown to be redundant, and phylogenetically distinct species can thrive in the same habitat and perform similar functions^7,45–47^. The taxonomic diversity and spatiotemporal heterogeneity of microorganisms, especially those of low-abundance and specialist species^48,49^, may hold information on processes shaping biodiversity beyond the scope of functional change^50^.

The study site Upper Lake Constance is a large deep perialpine lake shared by Germany, Switzerland and Austria (Fig. 1A). It has experienced various environmental perturbations since the end of the last glacial period across the Pleistocene-Holocene transition. These comprise climate change, regional floods, and also anthropogenic impact from human settlements leading to land modification and changes in the lake’s trophic state^51^. Like many lakes in the northern hemisphere^52,53^, Lake Constance became subject to eutrophication from increased household discharge, agricultural and industrial runoff in the mid-twentieth century but returned to an oligotrophic level in ∼2000 CE through wastewater purification initiated by the International Water Protection Commission for Lake Constance (IGKB)^54,55^. From a shotgun-sequenced sedimentary ancient DNA (*sed*aDNA) series, we identified bacteria and archaea with a reference-based approach, differentiated ancient taxa from potentially living microbes through time-associated DNA damage, and disentangled the dynamics of key groups responding to various environmental changes. We show that multiple factors, both of natural and anthropogenic origins, have contributed to the assembly of microbial communities since the Late Pleistocene, and that signals of dispersal and genetic changes might be observed among taxa displaying long-term persistence. These interwoven trajectories show that the system governing microbial communities in lake sediment is complex and its understanding can provide valuable insights in reconstructing paleoecological processes and long-term impact of humans on nature.

**Fig. 1.**
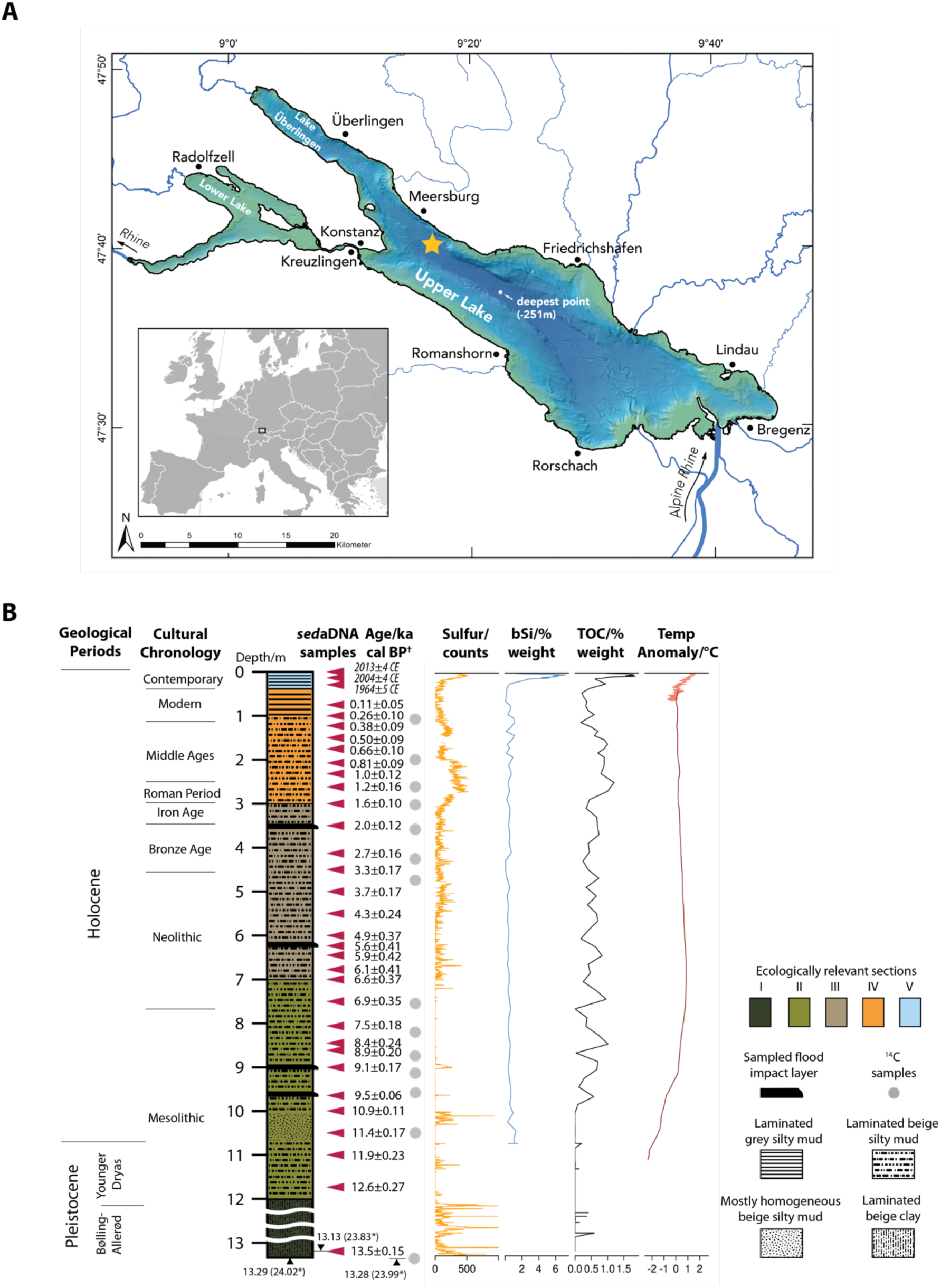
Drilling location and HIBO19 core diagram. (**A**) Map of Lake Constance. The star symbol shows the drilling location (UTM-32T, 521535, 5279467) of the HIBO19 sediment core. (**B**) HIBO19 core diagram (event corrected) with ecologically relevant sections described in Methods. Sections of high energy flood deposits (lithotype event 7 (LE7)^56^) were considered to have been deposited within days, therefore were removed from age-depth modelling and not shown in the core diagram. Depths of sediment below the removed sections were shifted up accordingly. Asterisk (*) indicates depth in the original core before shifting. Dagger (†) indicates unit of age unless noted otherwise. Sulfur, biogenic silica (bSi) and total organic carbon (TOC) contents are from Schaller et al. (2022)^56^. Temperature anomalies are from Kaufman et al., (2020)^57^ (0 to 12,000 ka cal BP) and NCEI NOAA (1850 CE until present) (Methods).

## Results

### Age-depth model and core chronology

The age of the HIBO19 core (Fig. 1B) spans from ∼13.5 ka cal BP (thousand calibrated years before the present) to ∼2014 CE (Common Era) as shown in the age-depth model (Fig. S1). The relationship between age and depth is approximately linear (excluding high-energy flood deposits). The higher sedimentation rate over the top 50 cm can be related to local hydrological activity, eutrophication and lower compaction near the sediment surface. The sampled core spans part of the Bølling-Allerød interstadial period, the Younger Dryas and the entire Holocene.

### Signatures of time-associated damage in sedimentary DNA

After DNA extraction and library preparation, the size distribution of selected individual *sed*aDNA samples, as well as the size distribution of the pooled library were measured (Data S1, Supplementary File). The individual libraries and the pooled *sed*aDNA library had highest molarity at 167∼183 bp. Taking the pooled library as an example (Supplementary File, “pool”), since indexing primers and adapters had 131 bp in length, the sizes of *sed*aDNA fragments therefore peaked at ∼38 bp, had an average length of 74 bp with 91% reads shorter than 200 bp. Individual sample libraries had similar size distribution. Shotgun sequencing of dual-indexed illumina libraries yielded between 8.9 and 77.2 (median 60) million clusters or single DNA reads per library for the 34 sediment libraries and 5.5 to 481.4 thousand (median 194.2) clusters per library for the ten control libraries (Table S2). After adaptor trimming, quality control, removal of low-complexity and duplicated reads, we obtained between 4.4 and 42.9 million merged reads per sample library (median 36.2) and between 1.7 and 99.5 thousand merged reads (median 26.8) per control library (Table 1). Among them, 0.5% to 2.7% were mapped to a bacteria or archaea reference genome in the GTDB r207 database (Table 1). These amount to a total of 6799 bacteria and 281 archaea species and are hereafter considered the closest species representatives of the recovered taxa. DNA damage estimation was based on taxonomic profiling, and the derived trend showed increasing damage over time, from ∼ 0.05 in the most recent sample to 2.0 to the oldest sample in the Holocene (Fig. 2).

**Fig. 2.**
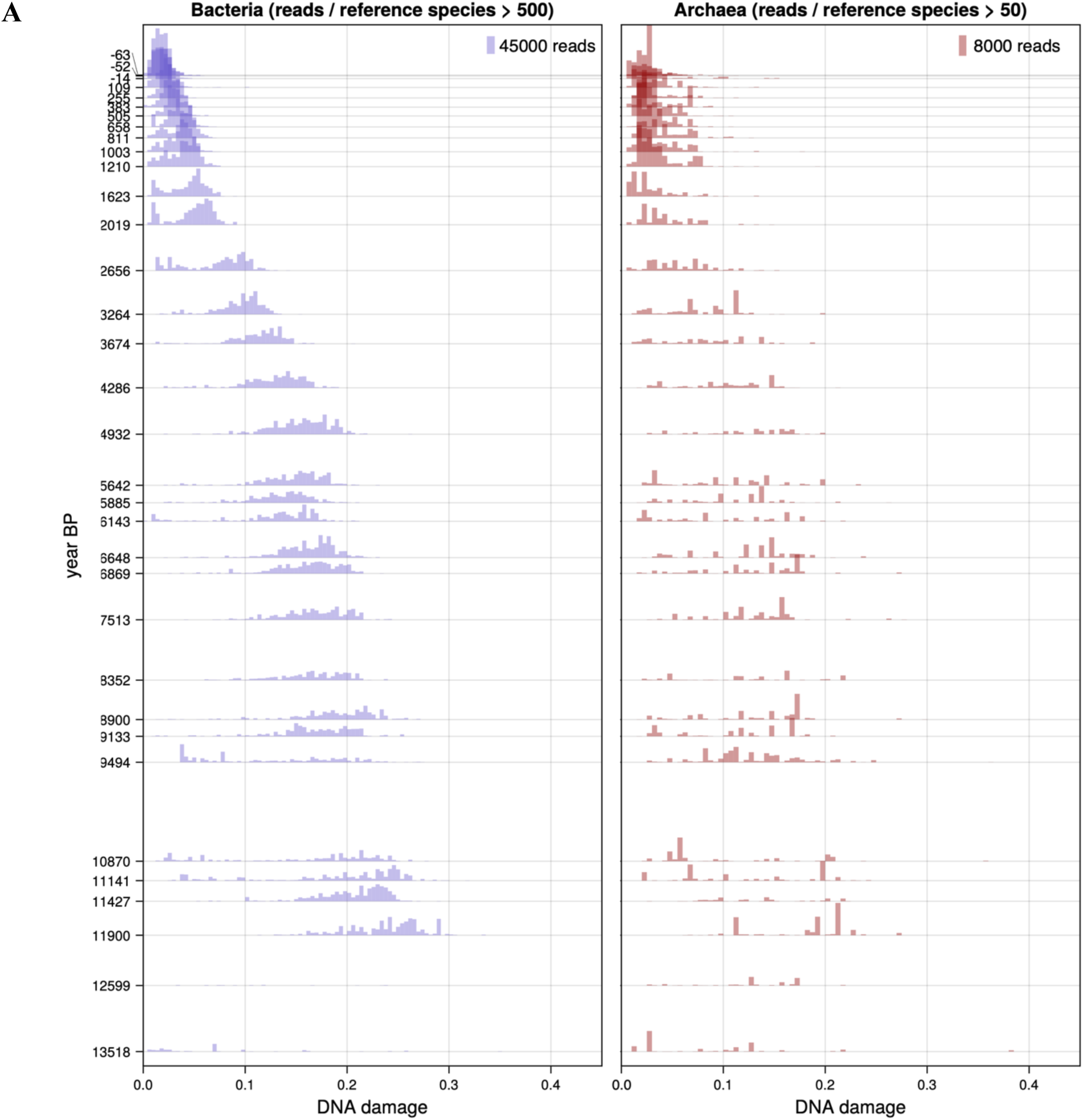
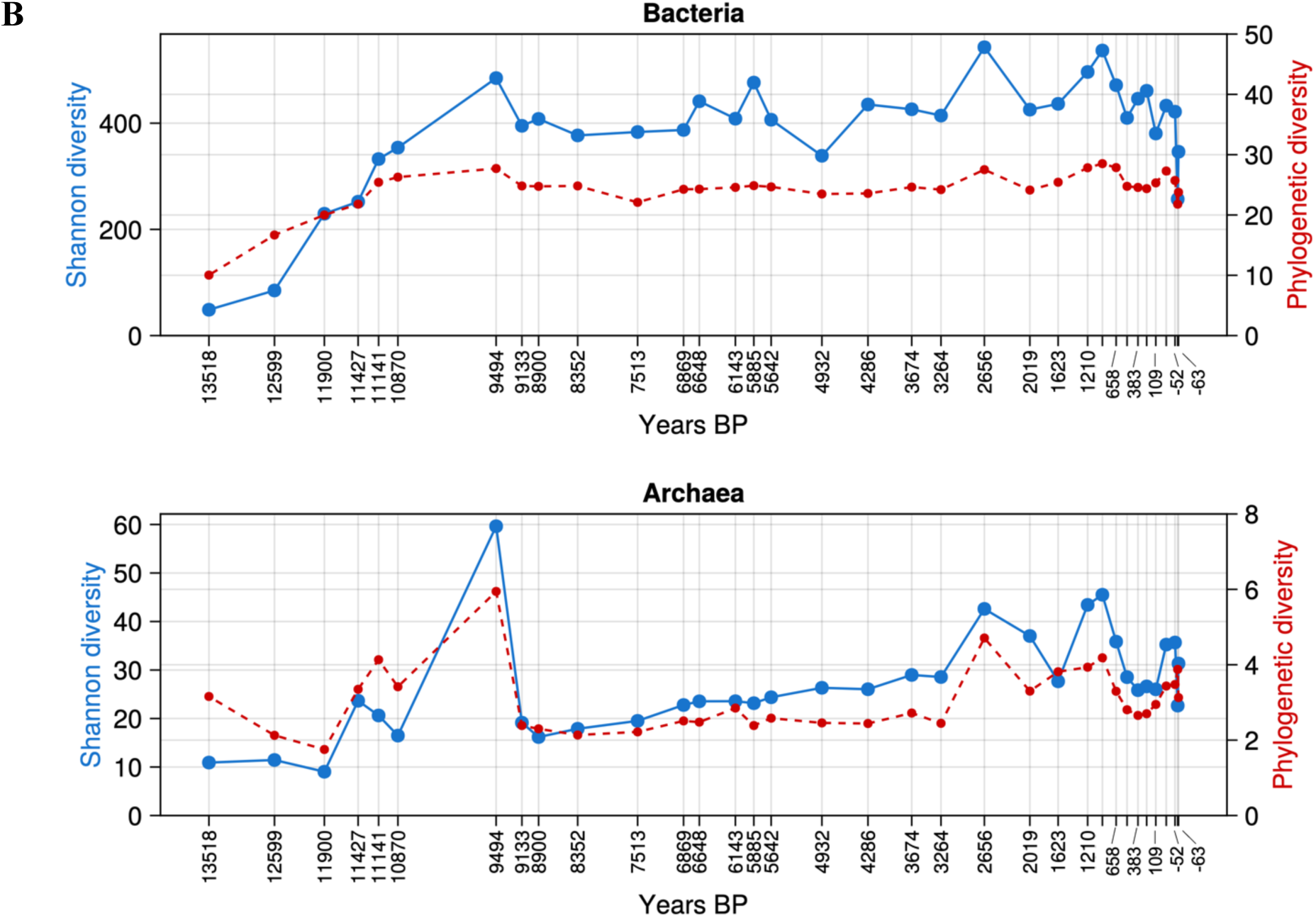
Overall diversity and DNA damage of bacteria and archaea. (**A**) DNA damage of bacteria and archaea. In each sample, reference bacterial species mapped by more than 500 reads and archaeal species mapped by more than 50 reads are included in the histogram. Bars are weighted by the read number assigned to each reference species. (**B**) Alpha diversity and phylogenetic diversity of bacteria and archaea. Reference bacterial species included in diversity index calculation have more than 100 reads per sample.

**Table 1.**
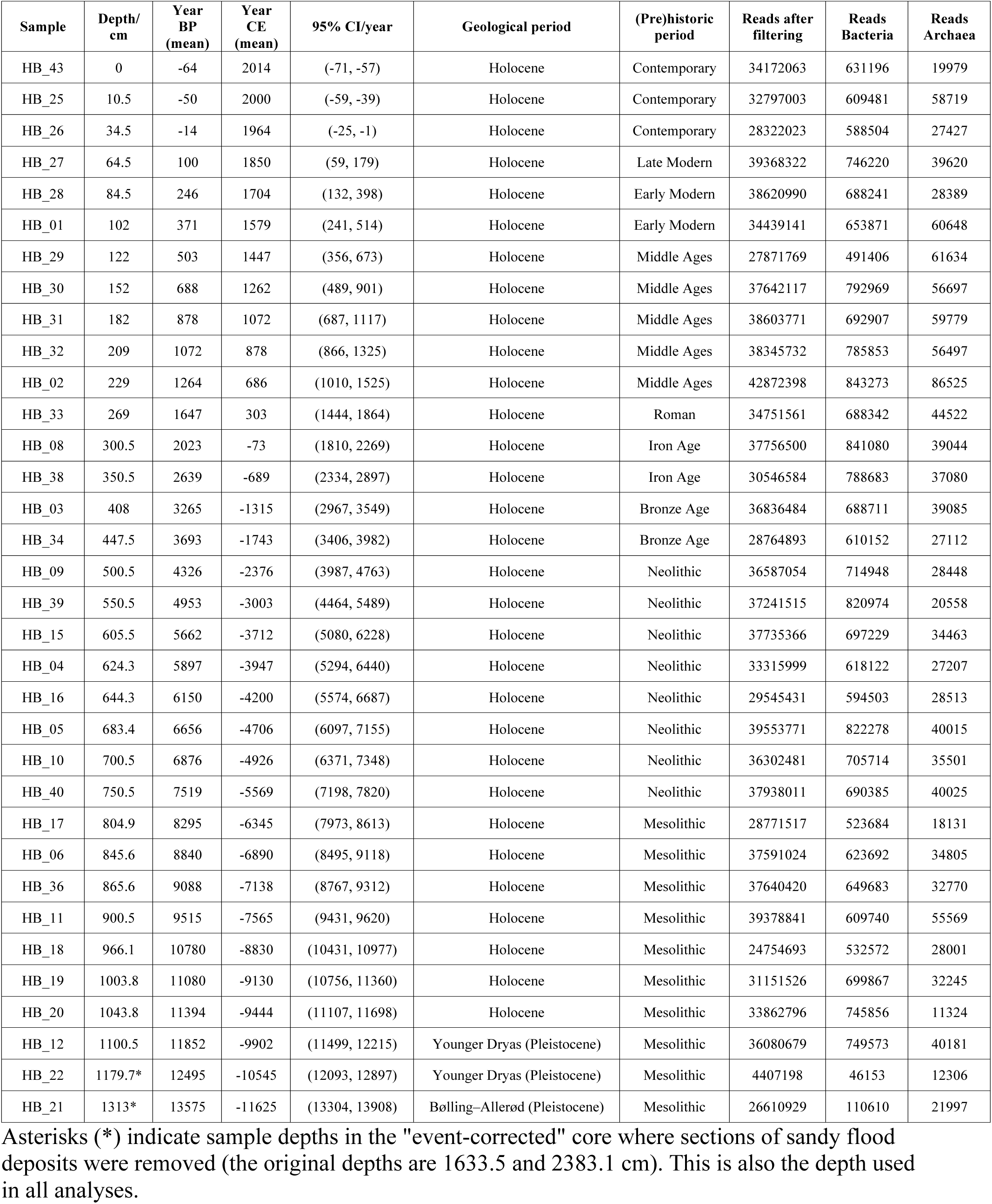
Sample metadata.

| Sample | Depth/<br>cm | Year<br>BP<br>(mean) | Year<br>CE<br>(mean) | 95% CI/year | Geological period | (Pre)historic<br>period | Reads after<br>filtering | Reads<br>Bacteria | Reads<br>Archaea |
| --- | --- | --- | --- | --- | --- | --- | --- | --- | --- |
| HB_43 | 0 | -64 | 2014 | (-71, -57) | Holocene | Contemporary | 34172063 | 631196 | 19979 |
| HB_25 | 10.5 | -50 | 2000 | (-59, -39) | Holocene | Contemporary | 32797003 | 609481 | 58719 |
| HB_26 | 34.5 | -14 | 1964 | (-25, -1) | Holocene | Contemporary | 28322023 | 588504 | 27427 |
| HB_27 | 64.5 | 100 | 1850 | (59, 179) | Holocene | Late Modern | 39368322 | 746220 | 39620 |
| HB_28 | 84.5 | 246 | 1704 | (132, 398) | Holocene | Early Modern | 38620990 | 688241 | 28389 |
| HB_01 | 102 | 371 | 1579 | (241, 514) | Holocene | Early Modern | 34439141 | 653871 | 60648 |
| HB_29 | 122 | 503 | 1447 | (356, 673) | Holocene | Middle Ages | 27871769 | 491406 | 61634 |
| HB_30 | 152 | 688 | 1262 | (489, 901) | Holocene | Middle Ages | 37642117 | 792969 | 56697 |
| HB_31 | 182 | 878 | 1072 | (687, 1117) | Holocene | Middle Ages | 38603771 | 692907 | 59779 |
| HB_32 | 209 | 1072 | 878 | (866, 1325) | Holocene | Middle Ages | 38345732 | 785853 | 56497 |
| HB_02 | 229 | 1264 | 686 | (1010, 1525) | Holocene | Middle Ages | 42872398 | 843273 | 86525 |
| HB_33 | 269 | 1647 | 303 | (1444, 1864) | Holocene | Roman | 34751561 | 688342 | 44522 |
| HB_08 | 300.5 | 2023 | -73 | (1810, 2269) | Holocene | Iron Age | 37756500 | 841080 | 39044 |
| HB_38 | 350.5 | 2639 | -689 | (2334, 2897) | Holocene | Iron Age | 30546584 | 788683 | 37080 |
| HB_03 | 408 | 3265 | -1315 | (2967, 3549) | Holocene | Bronze Age | 36836484 | 688711 | 39085 |
| HB_34 | 447.5 | 3693 | -1743 | (3406, 3982) | Holocene | Bronze Age | 28764893 | 610152 | 27112 |
| HB_09 | 500.5 | 4326 | -2376 | (3987, 4763) | Holocene | Neolithic | 36587054 | 714948 | 28448 |
| HB_39 | 550.5 | 4953 | -3003 | (4464, 5489) | Holocene | Neolithic | 37241515 | 820974 | 20558 |
| HB_15 | 605.5 | 5662 | -3712 | (5080, 6228) | Holocene | Neolithic | 37735366 | 697229 | 34463 |
| HB_04 | 624.3 | 5897 | -3947 | (5294, 6440) | Holocene | Neolithic | 33315999 | 618122 | 27207 |
| HB_16 | 644.3 | 6150 | -4200 | (5574, 6687) | Holocene | Neolithic | 29545431 | 594503 | 28513 |
| HB_05 | 683.4 | 6656 | -4706 | (6097, 7155) | Holocene | Neolithic | 39553771 | 822278 | 40015 |
| HB_10 | 700.5 | 6876 | -4926 | (6371, 7348) | Holocene | Neolithic | 36302481 | 705714 | 35501 |
| HB_40 | 750.5 | 7519 | -5569 | (7198, 7820) | Holocene | Neolithic | 37938011 | 690385 | 40025 |
| HB_17 | 804.9 | 8295 | -6345 | (7973, 8613) | Holocene | Mesolithic | 28771517 | 523684 | 18131 |
| HB_06 | 845.6 | 8840 | -6890 | (8495, 9118) | Holocene | Mesolithic | 37591024 | 623692 | 34805 |
| HB_36 | 865.6 | 9088 | -7138 | (8767, 9312) | Holocene | Mesolithic | 37640420 | 649683 | 32770 |
| HB_11 | 900.5 | 9515 | -7565 | (9431, 9620) | Holocene | Mesolithic | 39378841 | 609740 | 55569 |
| HB_18 | 966.1 | 10780 | -8830 | (10431, 10977) | Holocene | Mesolithic | 24754693 | 532572 | 28001 |
| HB_19 | 1003.8 | 11080 | -9130 | (10756, 11360) | Holocene | Mesolithic | 31151526 | 699867 | 32245 |
| HB_20 | 1043.8 | 11394 | -9444 | (11107, 11698) | Holocene | Mesolithic | 33862796 | 745856 | 11324 |
| HB_12 | 1100.5 | 11852 | -9902 | (11499, 12215) | Younger Dryas (Pleistocene) | Mesolithic | 36080679 | 749573 | 40181 |
| HB_22 | 1179.7* | 12495 | -10545 | (12093, 12897) | Younger Dryas (Pleistocene) | Mesolithic | 4407198 | 46153 | 12306 |
| HB_21 | 1313* | 13575 | -11625 | (13304, 13908) | Bølling–Allerød (Pleistocene) | Mesolithic | 26610929 | 110610 | 21997 |

Asterisks (\*) indicate sample depths in the "event-corrected" core where sections of sandy flood deposits were removed (the original depths are 1633.5 and 2383.1 cm). This is also the depth used in all analyses.

### Overall diversity and temporal distribution of bacteria and archaea

Our data feature the typical species abundance distribution observed in numerous macro- and microbial communities^58–60^ where most taxa are found in low abundances (e.g. almost 90% of taxa occurrences have <0.1% relative read abundance per sample). Additionally, most of these low-abundance taxa occur only in a few samples, and those that do persist longer in time tend to have high read counts (Fig. S2). This temporal heterogeneity of taxonomic composition is coupled with a structure in DNA damage that varies across both taxa and time. Most taxonomically assigned reads display DNA damage that increases with time, indicating that the cells died soon upon deposition into the sediment, hence forming a snapshot of communities at the time. Reads from a smaller number of taxa, in particular from the upper half of the core and from a few deep samples, show low DNA damage close to that observed in the youngest sediment, suggesting that these were microbes living in the sediment (Fig. 2A).

These two properties - the temporal structure of taxonomic composition and of DNA damage - are partially reflected by diversity metrics (Fig. 2B). Shannon (alpha) diversity and phylogenetic diversity showed similar trends of change for both bacteria and archaea. Diversities of bacteria increased during the four thousand years of the Pleistocene-Holocene transition and remained relatively stable throughout the Holocene. Diversities of archaea had an overall increasing trend and additionally showed higher values during ∼11.4 to 9.5 ka cal BP (Pleistocene-Holocene transition), ∼2.7 ka to 650 cal BP (Iron Age to the Middle Ages) and in the recent 150 years.

To reveal structure in this dataset of high taxonomic richness and temporal heterogeneity, we applied co-abundance network analyses on DNA sequence counts of non-singleton taxa in all samples (Fig. 3A). Their occurrences are seen strongly associated with changes in (pre)historic periods/time, temperature and sediment composition such as sulfur and biogenic silica (Fig. S3). Thirty-one percent of taxa in the network (1524 out of 4924), mostly low in DNA read abundance (Fig. S4), were closely connected communities. After clustering these 385 communities, we present them as seven major groups: 1) Long-term, stable taxa; 2) Long-term, unstable taxa; 3) Assemblages deposited by floods; 4) Uncommon taxa in early Holocene, 5) Uncommon taxa in late Holocene, 6) Microbes present since the Middle Ages (∼690 CE); and 7) Microbes present since the Modern time (∼1580 CE).

**Fig. 3.**
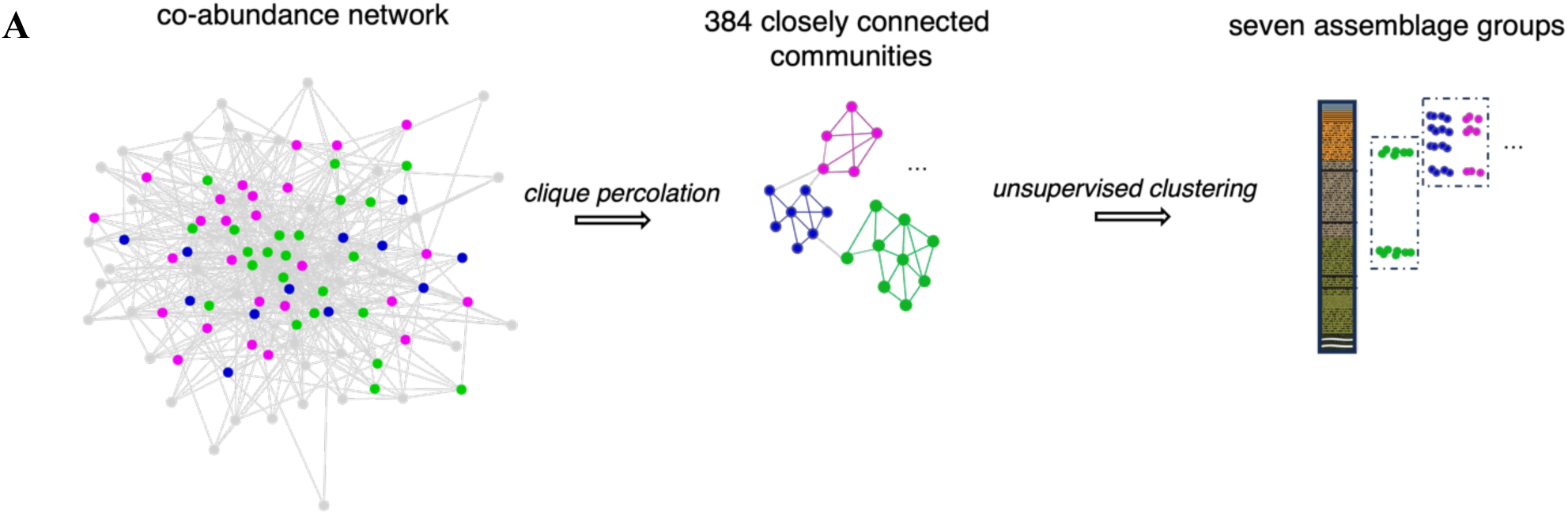
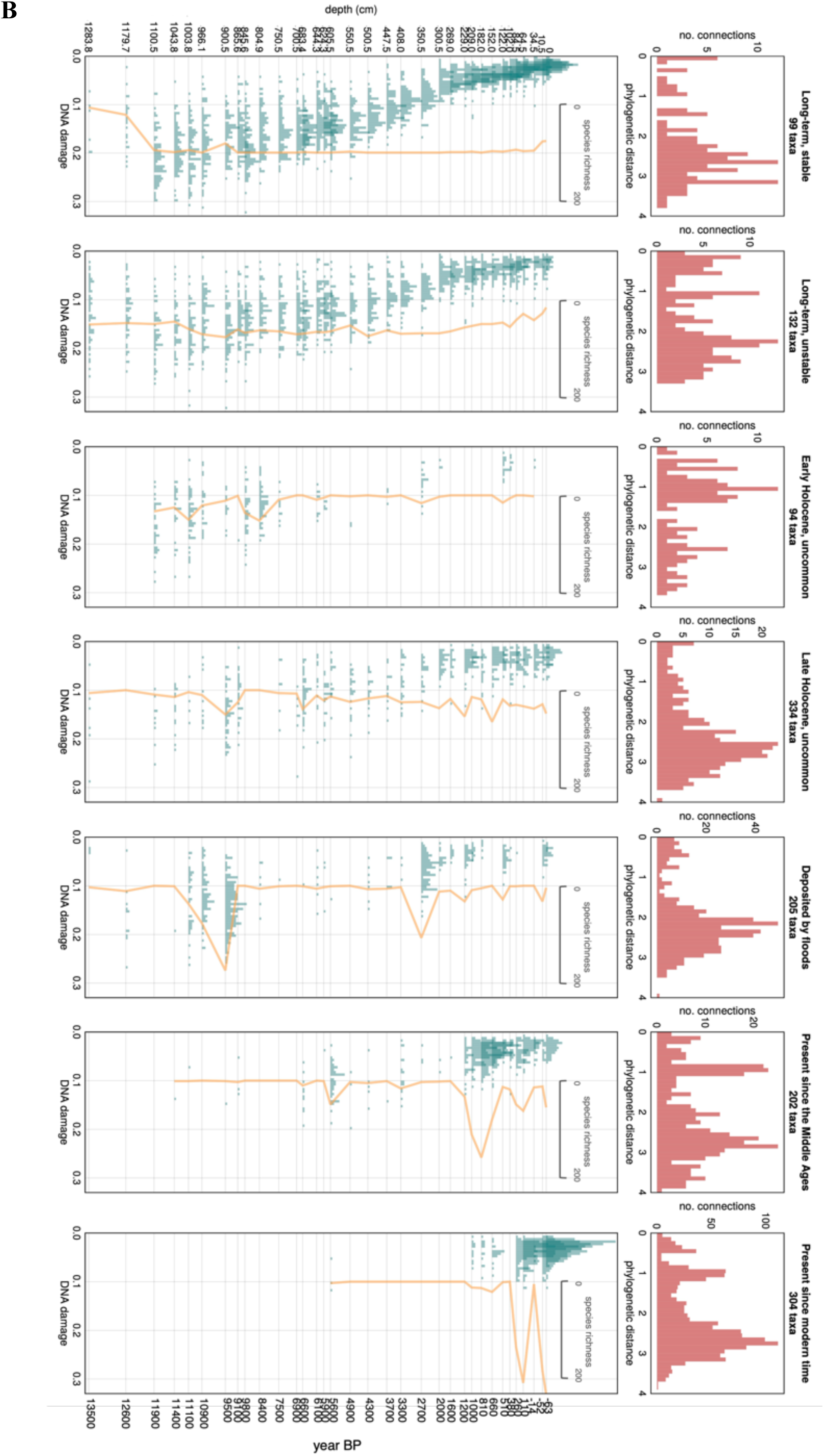
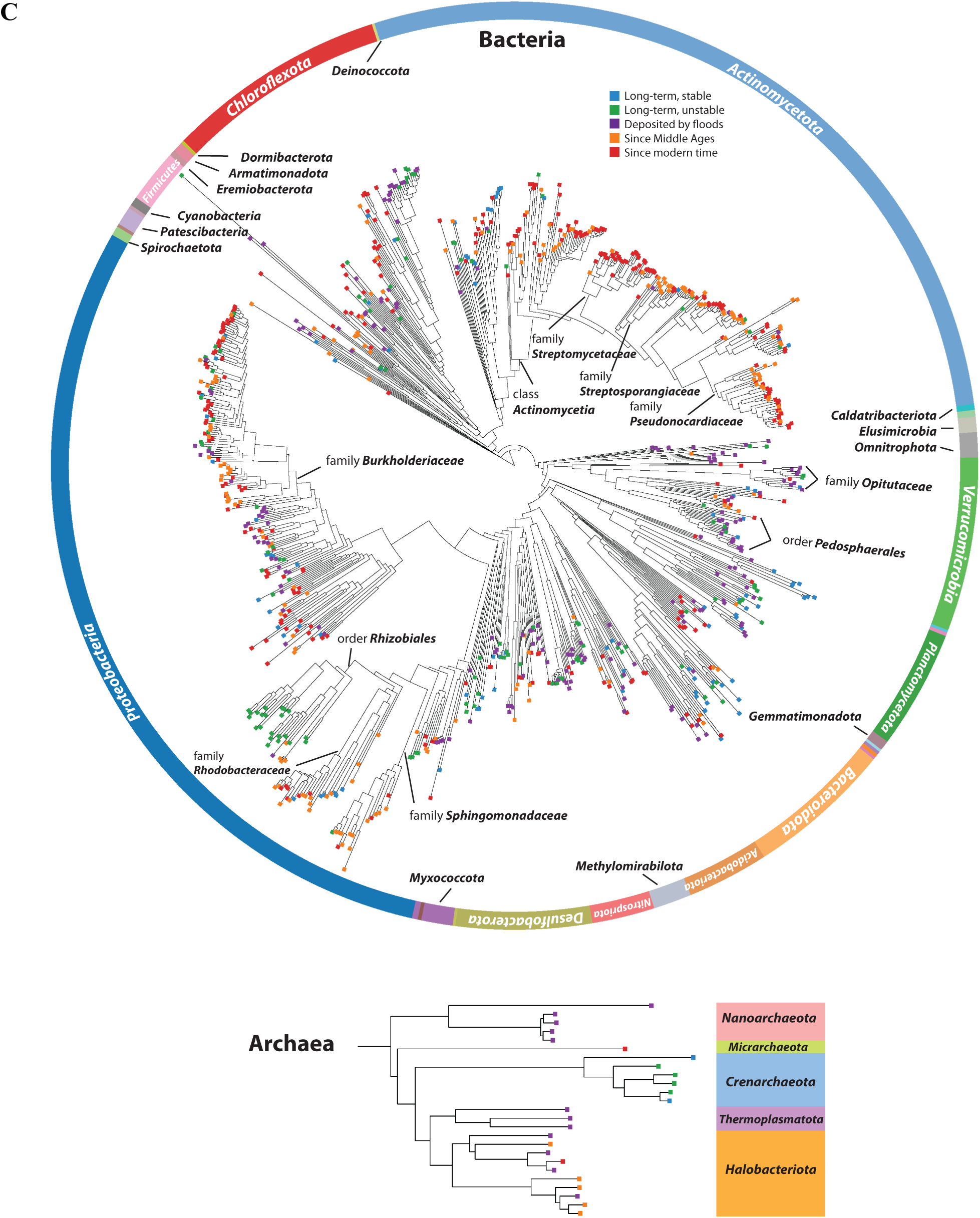
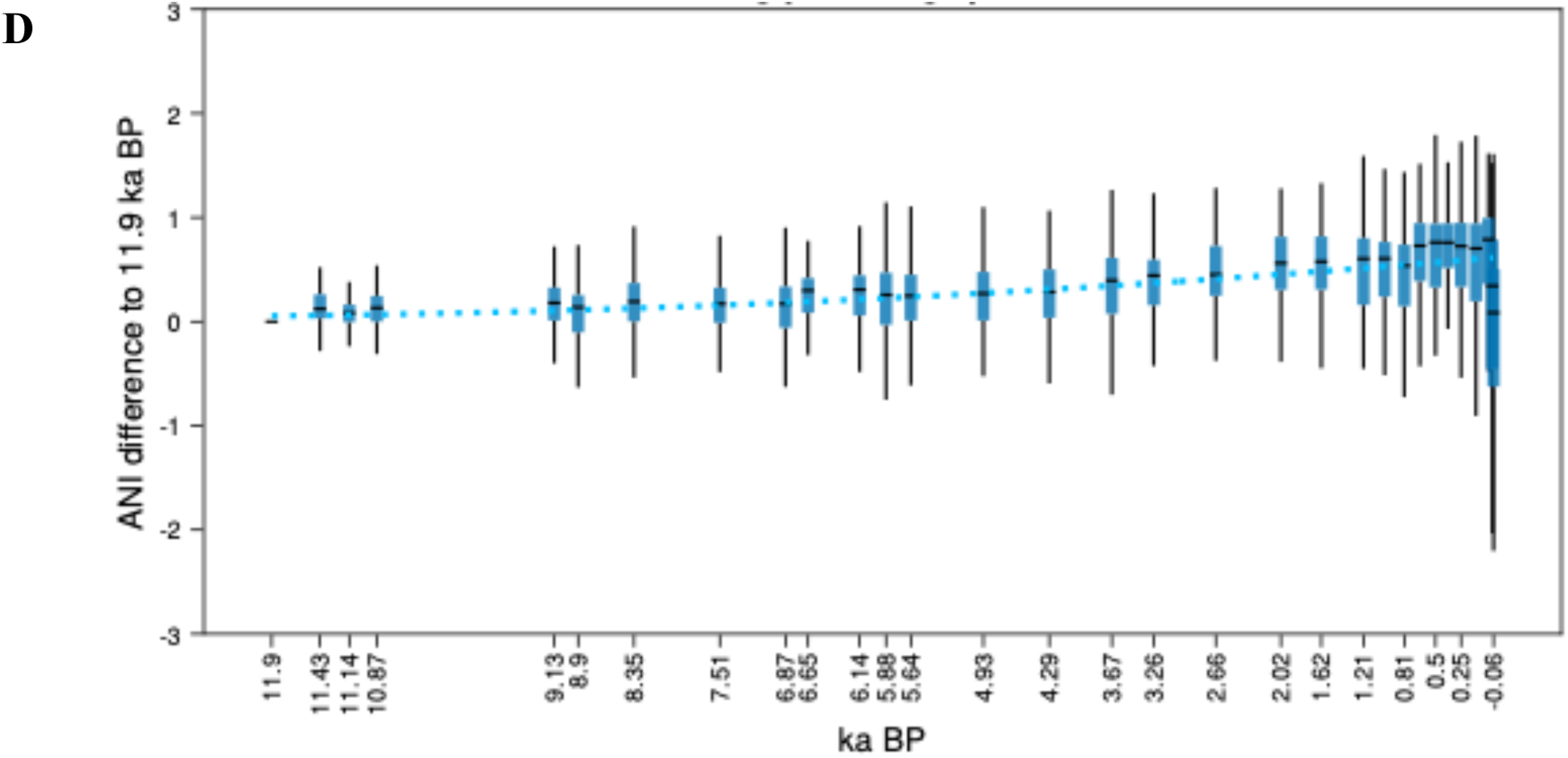
Occurrence pattern and diversity of seven major assemblage groups. (**A**) Workflow graphic of identifying communities in network and clustering into groups. (**B**) Assemblages are clustered into groups of similar occurrences. Upper panels show the phylogenetic distances of positively connected taxa in the network. Lower panels show the histogram of DNA damage of member taxa (teal bars) as well as taxonomic richness in each sample (orange lines). The tallest teal bar on this figure has 29 species, the shortest has one. (**C**) Taxonomic tree of bacteria and archaea, with tree tips coloured by group membership. Taxonomy is based on GTDB r207, and the tree was plotted using iTOL. The bacteria tree does not show the groups “Early-” and “Late-Holocene, uncommon”. The full bacteria tree showing all groups is in Fig. S5. (**D**) Changes in ANI difference - average nucleotide identity (ANI) relative to the values at 11.9 ka BP - for 97 selected long-term taxa. In each sample/time point, the box plot shows the median (dash in the blue box), interquartile range (i.e. 25%∼75%, blue box), and 1.5 times the interquartile range (whiskers) of ANI differences of these taxa. Dotted light blue line shows the fitted values from the longitudinal mixed model. ANI values from flood-impacted sediment samples and the top two samples (after the lake’s trophic state change) are excluded from the model.

### Long-term microbes show aspects of community stability

Two major assemblage groups represented communities that were long-term and predominant. The stable taxa among them (Fig. 3B, “Long-term, stable”) colonised the lake at the end of Younger Dryas and were reliably detected with high DNA abundance throughout the Holocene, many persisted through the lake’s late-20th-century eutrophication and re-oligotrophication. Their DNA damage indicated that most died after burial in the sediment. Being phylogenetically diverse, their type species were often found in various freshwater and soil environments. Bacterial taxa in this group included members in the order *Phycisphaerales* and family *Lumatobacteraceae* (Data S2). Some other taxa, although less persistent and abundant, were still detected over millennia (Fig. 3B “Long-term, unstable”). Members in this group are diverse, including e.g. anaerobic microbes in the genera *Methylomirabilis* (phylum *Methylomirabilota*) and *Sporomusa* (phylum *Bacillota*), bacterioplankton *Planktophila* (phylum *Actinobacteria*), and parasitic Phytoplasma (phylum *Mycoplasmatota*) (Data S2). This group additionally included taxa present primarily in the Pleistocene sediments (Fig. S6). Base on DNA damage, except for a few possibly ancient microbes such as those in the families *Haliangiaceae*, *Nitrospiraceae*, *Rhodocyclaceae*, and orders *Rhodospirillales* and *Anaerolineales*, many taxa in these sediments showed low DNA damage (<0.08), such as members in the genera *Labilithrix* and *Methyloceanibacter*, orders *Pirellulales*, *Anaerolineales* and *Sedimentisphaerales*, and phylum *Desulfobacterota* (Data S2), suggesting that they could have been living either due to their metabolic capacities in anaerobic environments or the presence of a subsurface aquifer^61^. For archaea in these two groups, detected taxa came exclusively from the order *Nitrososphaerales* (phylum *Thermoproteota*) (Fig. 3C).

Long-term microbes further offer a good opportunity for detecting signals of diversification and selection in unmanipulated environments. As an attempt to address this topic, we used mixed-effect models to estimate the temporal change of average nucleotide identity (ANI) for 97 well-identified (ANI > 0.95) long-term species between 11.9 ka cal BP to 1964 CE (Supplementary Text). The fitted curve showed increasing and slightly accelerating ANIs of approximately 0.5% per 10,000 years as a fixed effect, i.e. group characteristics (Fig. 3D). An abrupt increase of ANI starting from ∼1260 CE suggests the occurrence of lineages that are more similar to the reference genomes. A drop of ANI in the most recent two samples (not modelled) showed a fast divergence of detected taxa from the reference genomes, possibly suggesting a strong selection on new strains or species after the lake returned to an oligotrophic state.

Two other groups that contained uncommon taxa in the early Holocene or in the late Holocene (Fig. 3B, “Early-” and “Late-Holocene, uncommon”) were also identified through clustering. Overall, long-term taxa from these four groups were phylogenetically diverse but with little group-dependent structure. One exception was that taxa of the phylum *Planctomycetota* and the order *Rhizobiales* (nitrogen fixing bacteria, phylum *Proteobacteria*) mostly belonged to these long-term microbes (Fig. S5).

### Short-term assemblages reflect natural and anthropological impacts

Unsupervised clustering also revealed the occurrence patterns of short-term microbes that matched the lake’s geological events and stages of regional human history. A group of taxa primarily occurred in discrete sediment layers corresponding to flooding events in the lake’s history^56^ (Fig. 3B “Deposited by floods”). They were phylogenetically diverse and represented 22 phyla, 103 families and 136 genera. Abundant and prevalent taxa were seen in the families *Villigracilaceae* (class *Anaerolineae*, phylum *Chloroflexota*), *Nitrospiraceae* (phylum *Nitrospirota*), order *Elusimicrobiales*, and phylum *Desulfobacterota* (Data S5). Existing studies^62–64^ suggest that they may come from a diverse range of habitats including freshwater, soil and aquatic sediments.

The last two groups of microbes coincided with periods of fast development in human history: the Middle Ages (∼1.2 to 0.5 ka cal BP) and the Modern time (∼1700 CE) (Fig. 3B). Compared with earlier times, these two periods were associated with the prominent expansion of the bacterial class *Actinomycetia* (phylum *Actinomycetota*) and the family *Burkholderiaceae* (phylum *Proteobacteria*) (Fig. 3C). *Actinomycetia* is found in various habitats and has high bioactive potentials^65^. *Burkholderiaceae* is known to harbour saprophytic organisms, phytopathogens, opportunistic pathogens and primary pathogens for humans and animals^66^. In addition, the abundant DNA sequences peaking at lower DNA damage (<0.05) (Fig. 3B) suggested the prevalence of living microbes in the sulfur- and organic-rich sediments of these periods (Fig. 1B).

Specifically, high DNA abundance and taxonomic diversity of bacteria in the families *Streptosporangiaceae*, and *Pseudonocardiaceae* (phylum *Actinobacteriota*), *Rhodobacteraceae* and *Sphingomonadaceae* (phylum *Proteobacteria*) was observed since the Middle Ages (∼690 CE). Members of the family *Streptosporangiaceae* are mainly found in soil^67^, and members of *Pseudonocardiaceae* are known for their roles in environmental remediation and agriculture^68^. *Rhodobacteraceae* are mainly aquatic and *Sphingomonadaceae* are often isolated from soils, freshwater and plants^66^. The cyanobacteria genus *Synechococcus* also started to be prevalent during this period.

Archaeal taxa in this group belonged to the order *Methanomicrobiales* (methanogens), and all were likely to be living in the organic-rich sediments (DNA damage <0.06) (Data S6).

Microbes present since the Modern time showed the highest DNA abundance and taxonomic richness from ∼1700 CE until present, except for a plunge in these measures during the eutrophication phase (∼1960 CE) (Fig. 3B). Besides the families *Burkholderiaceae* and *Pseudonocardiaceae* that also dominated the previous group, abundant and prevalent taxa were seen in families *Streptomycetaceae* and *Nocardioidaceae* (phylum *Actinobacteriota*). Possible sources of these groups are soils, manure, plant hosts^67^ and industrial culture for their bioactive compounds e.g. antibiotics^69^. This group further included diverse taxa (38 families and 17 phyla) that appeared only recently, after the lake returned oligotrophic after four decades of anthropogenic eutrophication from ∼1960 to 2000 CE. Abundant ones were seen in the order *Desulfatiglandales*, family *Anaerohalosphaeraceae*, and genus *Arachnia* (family *Propionibacteriaceae*) (Data S7), possibly the sulfate-respiring and fermenting bacteria living in the sediment^70,71^. Members of these taxonomic groups have occurred in various habitats including freshwater and sediments^72^.

### Mapping qualities vary across assemblage groups

The mapping statistics of microbes were unevenly distributed across assemblage groups (Fig. 4). Taxa in the long-term groups generally had high DNA sequence abundance (Fig. S6) and also had overall higher mapping qualities to reference genomes, i.e. many had ANIs between 0.94 and 0.98 (Fig. 4A) and breadth ratios above 0.8 (Fig. 4B). Some taxa in the group “deposited by floods” also had higher mapping qualities. For the rest groups, most taxa had ANIs lower than 0.94 and breadth ratios below 0.8.

**Fig. 4.**
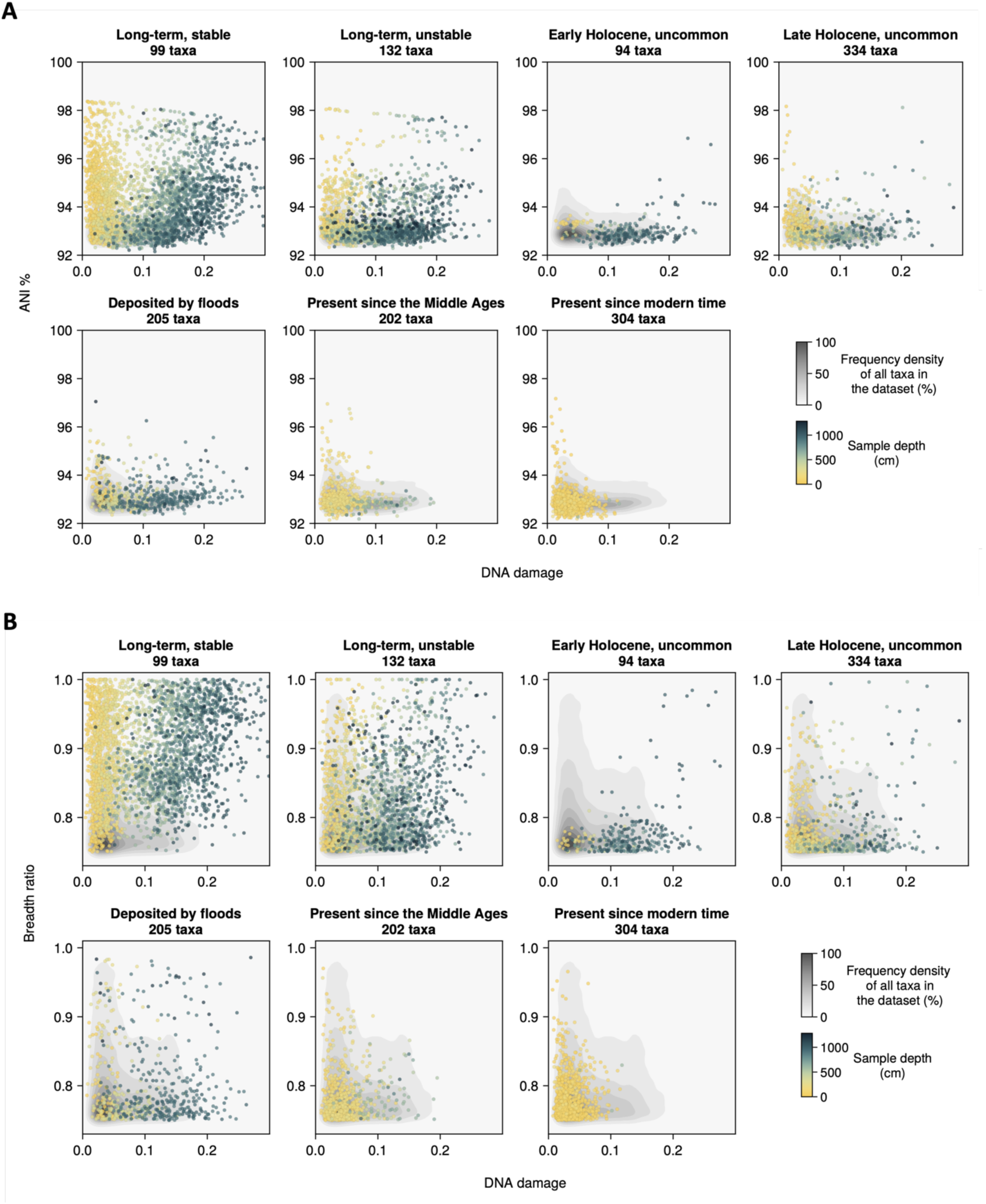
Mapping statistics of assemblage groups. (**A**) Mean average nucleotide identity (ANI) and (**B**) breadth ratio of sequences mapped to reference genomes, calculated at species level. Each panel represents taxa clustered into one assemblage group, each dot in a panel represents one occurrence of a taxon in a sample, color of the dot represents sample depth, and the shaded contour represents the density distribution of this metric for all taxa in the dataset.

## Discussion

Using ancient DNA, we show that prokaryotes in lake sediments document changes in the lake and its surrounding catchment at various scales and durations, including those related to climatic change, regional floods and historical development of human societies. Such changes are reflected by specific assemblages through their emergence and disappearance as well as variations in DNA read abundance. In line with studies on modern habitats^48,73,74^, our findings emphasize the role of rare taxa in reflecting environmental conditions^75^ but also reveal that this rarity, together with their tendency to occur within shorter times, could be the result of deposition from distant or specific origins (e.g. upstream, terrestrial, host organisms or human-sourced runoff) and short-term ecological or depositional events (e.g. floods and trophic state changes). Additionally, low-abundance microbes tend to show lower similarity to reference genomes than 95%, a commonly used threshold for identifying a prokaryote population, suggesting the existence of genetically more diverse populations than those represented by the reference genomes^76^. Although increasingly used in modern microbial metagenomics, metagenome-assembled genome (MAG)-based methods would not have recovered this diversity. Usually, 30 to 60 million paired-end reads (10 to 20 Gb of short-read data) are required to assemble several high-quality MAGs^77^. Since our dataset has an average per-sample sequencing depth of 33 million reads after filtration (Table S4), the assembled MAGs would only come from the most abundant organisms.

As our study is based on taxonomic profiling rather than de novo assembly of MAGs, the quality of taxonomic identification is highly dependent on the quality of reference databases. Therefore misidentification, especially of low-abundance and ancient microbes, may create noise or false temporal distributions of species that undermines network analyses. This limitation can be mitigated by applying integrated methods that combine reference-based and MAG-based taxonomic profiling^78^. Besides misidentification, long sampling time-intervals can further reduce the power of inference for long-term trends, and some short-term dynamics may not be picked up at all. This can be especially detrimental to analyses in the older time periods, as a decline in temporal resolution back through time is present in environmental metadata records (e.g. temperature and land use) as well. Our network inferences might therefore favor more recent times, and these limitations might have overall caused the loss of explaining power (i.e. decrease of degrees) of environmental variables in conditional networks, and that only a small proportion of species are identified as communities.

Studies that primarily target the molecular remains of living microbes in subsurface environments have concluded that there is only a weak association between their dynamics and paleoenvironmental proxies^22,79^. Indeed, the disjunct distributions of the potentially living microbes in the sediment of Lake Constance and other studies^19,31,32,80^ all point to a mixture of causes that shaped the active subsurface communities, such as geochemical composition changes (external, natural or anthropogenic causes) and adaptive succession of rare microbes (internal). However, examining beyond the active communities proves to bring more insight. Ancient microbial communities, as we demonstrate, can be indicators of paleoecological events beyond the modern timeframe^81^. For example, the recurrences of certain microbes (Fig. 3B “Deposited by floods”) corroborate flood events indicated by the lake’s sedimentological profile^56^ and are also congruent with regional flooding frequencies in the Alpine^82^. Also hinted at by Han et al^19^., this property of prokaryote communities complements that of single lineages^83^, functional genes^84,85^ or their metabolic products^84,86^.

In using ANI as a proxy for genetic changes of long-term microbes, we attempt a preliminary step towards characterizing genomic changes and potential evolution in environmental paleomicrobiology^17^. By tracing changes in the similarity to modern reference genomes, we show that the type strains of certain long-term taxa around Lake Constance were probably already selected for in the late Middle Ages (∼1260 CE). In recent decades, the fast deviation to reference genomes among these taxa, accompanied by local extinction of other long-term taxa also indicates that humans may be actively promoting diversification of prokaryotes. These dynamics at genomic level may precede or underlie changes in the presence and abundance of taxa. Anthropogenically induced evolutionary change is acknowledged as a consequence of recent impact^87^, and our data suggest that this extends to prokaryotes and encompasses not just directional selection. Tracing ecological and evolutionary dynamics through DNA from paleoecological archives will be highly consequential for predicting future biodiversity across the tree of life.

## Materials and Methods

### Coring campaign and geochemical data

Sediment drilling was carried out in Upper Lake Constance at a location 5 km east of Konstanz (UTM-32T, 521535, 5279467) from May to June 2019 during the Hipercorig drilling campaign^88^. In short, two sediment cores of 22 and 20 m in length were extracted in parallel using the Hipercorig hydraulic coring system at a water depth of 205 m. These two parallel cores were extracted as 1 to 1.5 m core segments, transported first on ice to the University of Konstanz, stored at 4°C in a cold room, and were later transported in an air-conditioned van to the University of Bern and stored in a 4°C cold room at the Institute of Geological Sciences. Storage at 4°C is similar to the sediment’s natural environment and was therefore considered sufficient. The two long cores and one short core, recovering the undisturbed sediment-water interface (length ∼1.5 m), were combined in a composite section representing the continuous sedimentary succession (core HIBO19; Schaller et al. (2022)^56^). Technical and operational details are published in Harms et al. (2021)^89^.

### Sediment dating and age-depth modelling

The sediment core was radiocarbon dated using plant remains, complemented by visual dating for the top ∼100 cm. We collected plant remains throughout the core by first collecting slices of bulk sediment (∼1 cm thick) at desired depths with a scalpel, washing the sediment chunks with distilled water and filtering through 190 µm disposable nylon sieves (E-D-Schnellsieb). Leaves and seeds of deciduous trees were then collected in glass vials and dried under a fume hood before being sent to Micadas (Alfred-Wegener-Institute, Bremerhaven) for radiocarbon dating^90^. A total of 16 samples with sufficient plant materials were sent for dating, and 10 dates were received. Dates collected in this study and in Schaller et al.^56^ for age-depth modelling are listed in Table S3. All radiocarbon dates were calibrated with IntCal20^92^. For the top-most ∼100 cm of the core where the dates were not older than 150 cal BP, we visually dated the core laminations by cross-referencing flooding events with historical records^91^. Age-depth model was built with the Bayesian Accumulation Chronology (BACON) implemented in the rbacon package (version 3.3.1)^93^ (Supplementary Text).

The (pre)historic period that each sample belongs to was inferred with reference to Rösch (2021^96^, Table 5) and from the cultural chronology in Tinner et al. (2003^97^, Table 2). Based on cultural chronology and the stratigraphic units and lithotypes defined in Schaller et al.^56^, we categorise the core’s ecologically relevant backgrounds and events into five chronological sections (Fig. 1B, indicated by core colours; Supplementary Text).

### Environmental metadata

Temperature data in the past 12000 years (0 to 12 ka cal BP) were represented by the 30-60° N median temperature anomalies relative to 1800-1900 CE from Kaufmann et al. (2020)^57^. Temperatures after 1850 CE were represented by the Northern Hemisphere January to December average land temperature anomalies (with respect to the 1901-2000 CE average) downloaded from the NCEI NOAA, (https://www.ncei.noaa.gov/access/monitoring/climate-at-a-glance/global/time-series) and were roughly adjusted to 1800-1900 CE by referring to the anomaly in 0 cal BP/1950 CE in the former temperature series.

### Sedimentary DNA extraction, library preparation and sequencing

In November 2019, around five months after drilling, core segments were halved along the height of the core liner with laser and steel plates. At each sampling depth, the surface layer of the sediment was removed with a single-use sterile surgical blade. Immediately afterwards, at the surface-removed area of the sediment, two to three sediment samples were collected, each by inserting a single-use, top-removed syringe into the sediment. Around 3 cm3 sediment was collected by each syringe and was then stored in a 6 mL tube. Tubes containing sediment samples were transported in a styrofoam box filled with dry ice to the University of Konstanz and stored there in a −80 °C freezer.

### *sed*aDNA extraction and library preparation were carried out in an ancient DNA

designated lab at University of Konstanz. We selected a set of 42 sediment samples and extracted *sed*aDNA using the method described in Murchie et al. (2021)^98^ with the modification of doubling the input sediment quantity, i.e. from 250 mg to 500 mg. Each extraction yielded 100 µL extract from around 500 mg of sediment. For each sample, 50 µL of DNA extract was used for library preparation.

Libraries for shotgun sequencing were prepared following the protocol from Meyer and Kircher (2010)^99^ with the modification of dual indexing. After the adapter fill-in step, library concentration was measured with 1 µL sample in digital droplet PCR (ddPCR) with the SYBR Green qPCR Master Mix kit (Thermo Fisher) performed in a standard molecular biology lab. The final indexing cycle was determined as the equivalent of Ct, where Ct is the number of cycles reaching the exponential growth phase in the ddPCR. After indexing PCR, libraries from all samples were subsequently quantified using Agilent High Sensitivity DNA Kit and the 2100 Bioanalyzer instrument. After examining the DNA size distribution plot, we excluded two libraries with a narrow distribution at short size, which indicated low DNA complexity and high duplication rate. Eventually 44 libraries, including 34 *sed*aDNA samples and 10 extraction and library preparation controls (1 µL each) were pooled equimolarly and cleaned up using the MagBio system.

The pooled library showed a fragment length distribution of 20∼200 bp, peaking at ∼40 bp (excluding 130 bp of adapter sequences; Fig. S2). The library was then diluted with HPLC water to 1 nM in 30 µL, stored in a safeseal tube and sent to Fasteris™, Switzerland for sequencing. Sequencing was run on a single lane of the S2 Flow Cell in the Novaseq 6000 System with paired-end 2 x 100 bp mode.

### Sequencing data processing and taxonomic profiling

Raw reads were delivered as fastq files. We processed raw reads by removing adapter sequences merging paired-end reads using fastp^100^. Low-complexity reads removal and deduplication were performed with SGA (dust-threshold=4)^101^. Duplication rates estimated as the proportion of removed reads through deduplication ranges from 0.30 to 0.03 (median 0.20). We then trimmed poly-N heads and tails and kept reads with a minimum length of 30 bp. In each processing step the numbers of removed reads were kept track in Table S4.

For taxonomic profiling of bacteria and archaea, we mapped all filtered reads against reference genomes in the GTDB r207^102^ database using the aMAW pipeline^13^. In this pipeline, reads were first mapped to reference genomes using bowtie2^103^, then taxa (as in bam files) were filtered based on breath ratio and coverage evenness etc. using bam-filter^104^ to remove spurious results. Most species-level mapping to the reference have average nucleotide identities (ANIs) of around 0.93 (Fig. S7) and only a small percentage of taxa pass the commonly used threshold 0.95 for species identification 0.95. This suggests that most identified taxa were in the same genera as the reference species, but were themselves not represented in the reference database. To preserve this observed taxonomic richness, we therefore chose to characterise taxa having at least 0.92 in ANIs. Output bam files from this step were finally used as input to metaDMG^105^ to find the lowest common ancestor (LCA) and to estimate DNA damage.

### DNA damage estimation and taxon filtration

The average DNA damage of each identified taxon was described by the D_fit, Z_fit and rho_Ac statistics from metaDMG output. D_fit (“damage”) is the fitted frequency of deamination at the first base of a DNA molecule, Z_fit (“significance”), defined as the ratio of D_fit to the standard deviation of the data, represents the certainty of D_fit being non-zero^105^. rho_Ac, defined as the correlation between damage estimate and baseline deamination rate represents the goodness of fit. Since DNA damage is a signature of age, the damage estimates (D_fit) of all identified taxa in a sample should theoretically have a distribution that is associated with the sample age. This signal would then be blended by noises due to poorly identified taxa (due to either inadequate reads or missing reference genome), or modern DNA contamination. For this study, we consider microbes that survived longer in the sediment after burial, or those that are still alive at the time of sampling are also an integral part of the microbiota-of-interest, hence we chose not to separate them from ancient taxa based on DNA damage. In fact, the duration microbes can persevere in the sediment might differ, and their DNA damage would have a more continuous distribution than exhibiting a cutting point that separates the living ones from ancient ones. Therefore, we applied loose filtering criteria where taxa with a minimum Z_fit of 0.3, a range of rho_fit of −0.6 to 0.6, and a minimum of 50 reads were kept. Data from the control libraries were filtered in the same way as samples except for a minimum read count of 10. Any species-level identifications that are present in the controls are thereafter filtered out from the samples.

### Diversity indices, taxonomic tree and phylogenetic distance

Shannon diversity was calculated using the hillnumbers (q=1) function in the Diversity.jl package, where species-level read counts were used as the abundance measure. Phylogenetic diversity was calculated using the meta_gamma (q=1) function from the Phylo.jl package operated on the metacommunity object. The metacommunity object was generated with the Metacommunity function, using species-level read counts and a taxonomic tree as input. Taxonomic trees of bacteria and archaea were built using phyloT^106^ based on GTDB r207 taxonomy, and annotated with iTOL^107^. These trees contain branch lengths and clade support values therefore calculation of taxa phylogenetic distance is possible. Phylogenetic distance between any two species was calculated using the distance function in the Phylo.jl package.

### Network analysis and identification of closely connected microbe assemblages

We built read co-abundance based networks to find structure of microbial communities (Supplementary Text). We then applied clique percolation^109^ to identify closely connected microbial assemblages in the network. Function clique_percolation from Graphs.jl package was applied to the conditional network, where clique size (k) was set to three. To further extract structure from the derived 385 communities (1532 taxa), we then applied spectral clustering to cluster these communities into seven groups (Supplementary Text). We assessed the importance of environmental variables such as temperature, sedimentological composition and (pre)historic periods in microbial communities by comparing the number of connections they have in the networks. To evaluate the longer-term influence of these environmental variables, we compared three networks, each containing taxa that appear in at least two, three, and five samples respectively. The number of connections of each environmental variable representing land use and (pre)historic periods was calculated with the degree function in Graphs.jl.

## Acknowledgments

The authors thank Antonio Fernandez-Guerra, Janko Tackmann, Peter A. Seeber, Anna Chagas and Alexandra Schmidt for advice and practical support. We also thank the Centre for Ancient Environmental Genomics at the Globe Institute, University of Copenhagen (Denmark) and the bwUniCluster 2.0 system of Baden-Württemberg’s Universities and Universities of Applied Sciences (Germany) for providing computation resources.

## Author contributions

Conceptualization: Y.W., L.S.E.

Methodology: Y.W., F.S.A., A.S., M.W.P., L.S.E.

Software: Y.W., M.W.P.

Validation: Y.W., L.S.E. Formal analysis: Y.W. Investigation: Y.W., E.M., S.S.

Resources: Y.W., D.S., M.W., F.S.A., L.S.E.

Data curation: Y.W.

Writing - original draft: Y.W.

Writing - review & editing: Y.W., D.S., E.M., M.W., S.S., F.S.A., A.S., M.W.P., L.S.E.

Visualization: Y.W., M.W.

Supervision: M.W.P., L.S.E. Project administration: L.S.E.

Funding acquisition: Y.W., A.S., L.S.E.

## Data and materials availability

Raw sequence data are deposited to the European Nucleotide Archive (ENA) under the project accession PRJEB76926. The age-depth model was built with rbacon^93^. Data analyses and plotting used Julia (version 1.9.4) and R (version 4.2.3). Custom code and files necessary to process data, run analyses and produce results are available at: https://github.com/wangyi91/HIBO19_sedaDNA_prok.

