## Supplementary Materials for "Microbial sedimentary DNA from a cultural landscape disentangles the impacts of humans and nature over the past 13.5 thousand years"

**This PDF file includes:**

Supplementary Text
References of Supplementary Text
Figs. S1 to S7
Tables S1 to S4
Supplementary File

**Other Supplementary Materials for this manuscript include the following:**

Data S1 to S7

### Supplementary Text

#### Core sections

We categorise the core's ecologically relevant backgrounds and events into five chronological sections (Fig. 1B, indicated by core colours): I. The oldest section of the core reaches back to the end of the Pleistocene, from middle Bølling-Allerød interstadial until early Younger Dryas (~13.8 to 12.5 ka cal BP, depth 2400 to 1200 cm). During this warm period, massive sand beds and sandy laminae were deposited likely by the active subaquatic channel from river Seefelder Aach in several events, with occasional high concentration of terrestrial plant remains in the fine sandy laminae. Organic carbon content in the lake during this period is very low. Two *sed*aDNA samples (depth 2383.1 and 1633.5 cm) in the dataset are from this period. II. Early Younger Dryas until the beginning of the Neolithic Age in the Holocene (~12.5 to 6.9 ka cal BP, depth 1200 to 700 cm). Although organic matter content remains low until 11.2 ka cal BP (depth ~1000 cm), elevated biogenic silica value during 12.5 to 11.9 ka cal BP (depth 1164 to 1104 cm) indicates an increased in-lake productivity during the Younger Dryas cold period. The most notable clastic event is the Flimser rockslide (~9500 cal BP, depth ~900 cm). Ten *sed*aDNA samples (depths from 1100.5 to 700.5 cm) are from this period. Among them, samples at depths 966.1 and 900.6 cm are at flood-deposited sediment layers based on lithotype analysis. III. Beginning of Neolithic Age until the end of Iron Age/beginning of Roman time (~6.9 to 2 ka cal BP, depth 700 to 300 cm). Sulfur content becomes elevated here from ~6.9 ka cal BP (depth 700 cm), coinciding with the start of the Neolithic age. Ten *sed*aDNA samples (depths from 683.4 to 300.5 cm) are from this period. Among them, samples at depths 624.3 and 350.5 cm are at flood-deposited sediment layers. IV. Roman time until the lake's major anthropogenic eutrophication (~2 ka cal BP to 1960s CE, depth 300 to 35 cm). Sulfur content increases dramatically during the Roman time and early Middle Ages (~2 to 1 ka cal BP) and contemporary time (after ~1985 CE). Eleven *sed*aDNA samples (depth 300.5 to 34.5 cm) in the dataset are from this period. V. The lake after returning to oligotrophic state (after 2000 CE). Two *sed*aDNA samples (depth 10.5 and 0 cm) are from this period.

#### Age-depth model

We built the age-depth model with the Bayesian Accumulation Chronology (BACON) implemented in the rbacon package (version 3.3.1)[1] with a total of 27 dates: 10 radiocarbon dates collected in this study, five published radiocarbon dates collected from HIBO19 core[2] (Table S3), one published radiocarbon date[3] on the Flims rockslide event[4], whose impact layer is visible in the HIBO19 core, and 11 visual dates based on floods occurring along the upper Rhine valley during the Pleistocene-Holocene transition period between 13.5 and 12 ka cal BP[2], the core has a large amount of sand deposition from 12 m and downwards, which disrupted the age-depth relationship. To mend this and make age estimation below the flood deposits possible, we created an "event-corrected core" by excluding these sandy deposits (depths of affected sections were shifted upward, accordingly). Dates and depths of this event-corrected core were used as input of the age-depth model. Ages between any two adjacent dated depths and corresponding uncertainties were linearly interpolated.

### Co-abundance network

As sampling time intervals in this dataset range from decades to centuries, potentially longer than the generation times of active prokaryotes, we assumed that the temporal association of sequentially assembled microbial communities can be ignored, and the co-occurrence of taxa among samples are mainly explained by environmental factors. Based on this reasoning, we interpreted the bioinformatic communities identified in the networks as reflecting the dynamics of ecological conditions that supported the survival of these microbes.

Networks were constructed on 4924 bacterial and archaeal taxa occurring in at least two samples with the learn_network function (sensitive=true, heterogeneous=false) in the FlashWeave.jl package[5]. We used both k=1 for constructing a conditional network that includes only direct associations between nodes, and k=0 for constructing a univariate network that includes both direct and indirect associations. To evaluate the effect of environmental factors, we further included the following environmental meta-variables (MVs): sulfur, biogenic silica data[2], continuous time (cal BP), 30-60° N median temperature anomalies and land use data (restricted to the approximate catchment area of Lake Constance) (Methods: Retrieval of environmental metadata). MV value at each sampling depth was estimated by linear interpolation between the nearest two data points from the original data source. MVs that have absolute values were standardised while those in percentage were used as is.

### Spectral clustering of bioinformatically derived communities

To extract structure from the 385 bioinformatically-derived communities (1532 taxa), we applied spectral clustering to cluster these communities into five to eight groups. Each community was represented by a vector of numbers of detected taxa for all samples. To mitigate stochasticity of clustering, k-mean clustering on eigenvectors was run for 1000 times and a consensus result was obtained by requiring two communities being connected for at least 900 times (i.e. frequency $\geqslant$ 0.9). In the consensus result, the top seven community groups contained 89.4% of taxa (1370 out of 1532), a result that was insensitive to how many clusters were initially specified. These seven groups were hereafter named based on the geological activities and historic periods that they corresponded to: 1) Long-term, stable taxa; 2) Long-term, less stable taxa; 3) Uncommon taxa in the early Holocene; 4) Uncommon taxa in the late Holocene; 5) Assemblages deposited by floods: their read counts peaked in depths where the sediments are known to be flood deposits: 966.1, 900.5, 624.3 and 350.5 cm. The sample at 966.1 cm is located directly above the medium-to-high energy deposits of the underwater channel. The sample at 900.5 cm sits at the impact layer of the Flimser Rockslide, and the samples at 624.3 and 350.5 cm is located exactly at a flood-impacted layer; 6) Microbes present since the Middle Ages (~690 CE); 7) Microbes present since modern time (~1580 CE).

### Longitudinal analysis of ANI for long-term taxa

To find signals of potential intraspecies diversification and selection, we modelled the change of average nucleotide identity (ANI) over time for long-term taxa using mixed-effect models (Supplementary Text). Among taxa that were identified as the same species to the reference genome, we retrospectively expected their similarity to their modern references to increase overtime throughout the Holocene. Under the combined effects of evolutionary change in a natural, stable environment and DNA damage that increases approximately linearly with time (Fig. 2A), ANI should increase roughly linear with time.

Taxa that had at least 95% ANI in all samples (excluding those at depths 966.1, 900.5 and 350.5 cm as they were under major flood impact) were included. We built mixed-effect models which allow a fixed effect of time as group characteristics and random effects on individual species. Note that rates of evolutionary change among prokaryotes can vary by three orders of magnitude[6], the estimate of fixed effect might mainly account for the effect of DNA damage, and ANI differences caused by evolution were mainly reflected in the random effects of individual species. Models included samples from depth 1100.5 cm (11.9 ka cal BP) until 34.5 cm (1964 CE) excluding the Flimser Rockslide sample at 900.5 cm. The Flimser Rockslide sample was excluded because its taxa composition did not represent the same environment as the other samples. Samples from the Pleistocene and the most recent two samples (from after the lake returns to oligotrophic state following anthropogenic eutrophication) were excluded because communities were very different in those time periods hence were considered not to meet the assumption of the models. A total of 97 species were included in the models. Mixed-effects models were built using the package MixedModels.jl with quadratic terms of time included to model the nonlinear effect of time, when there is a better fit (smaller AIC). The model can be written in Julia's syntax as:

$$\text{ANI} \sim 1 + \text{time} + (1 + \text{time} \mid \text{species}) \tag{1}$$

And with quadratic terms:

$$\text{ANI} \sim 1 + \text{time} + \text{time}^2 + (1 + \text{time} + \text{time}^2 \mid \text{species}) \tag{2}$$

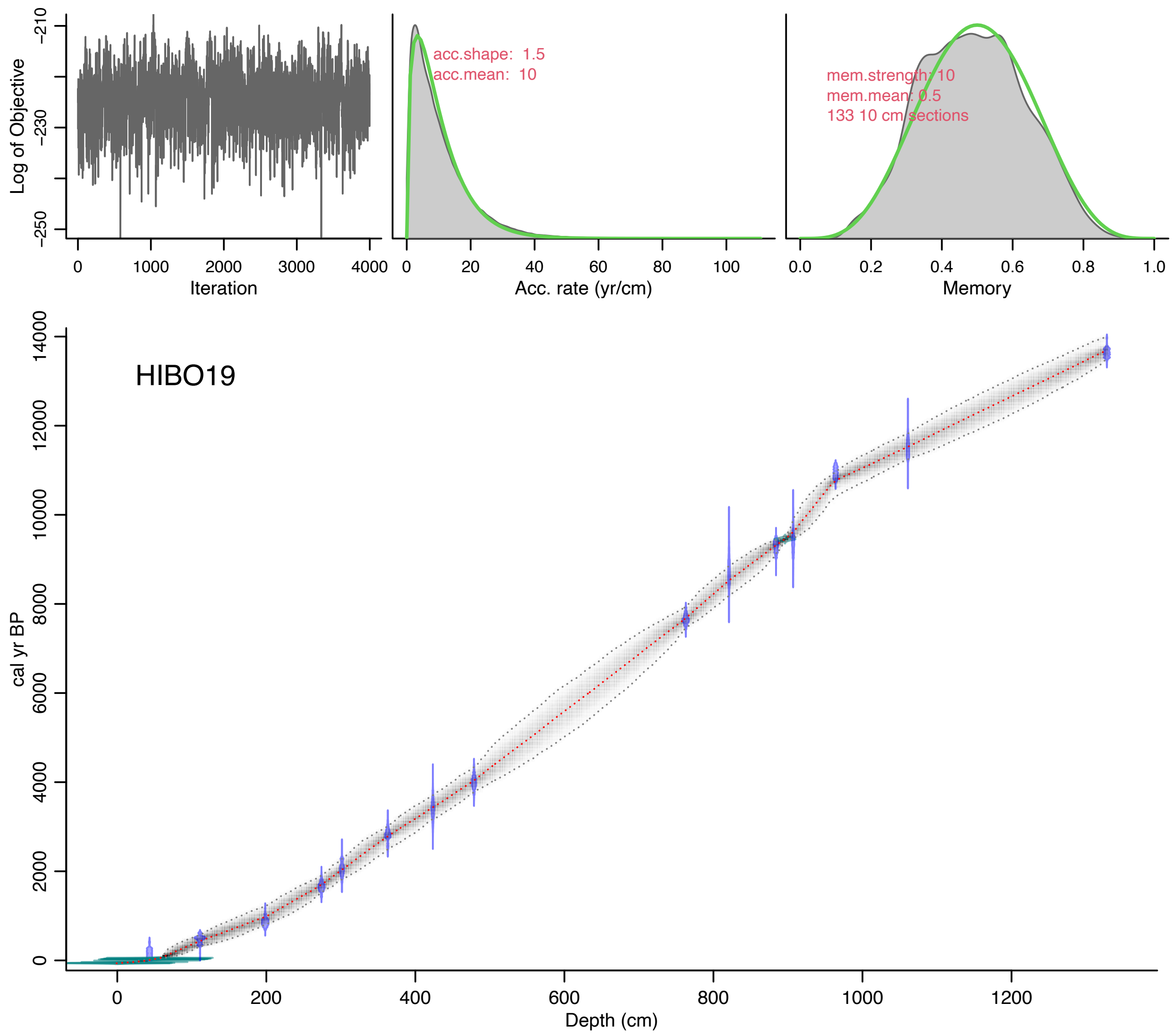


**Fig. S1.**
Age-depth model of the HIBO19 core (event-corrected).

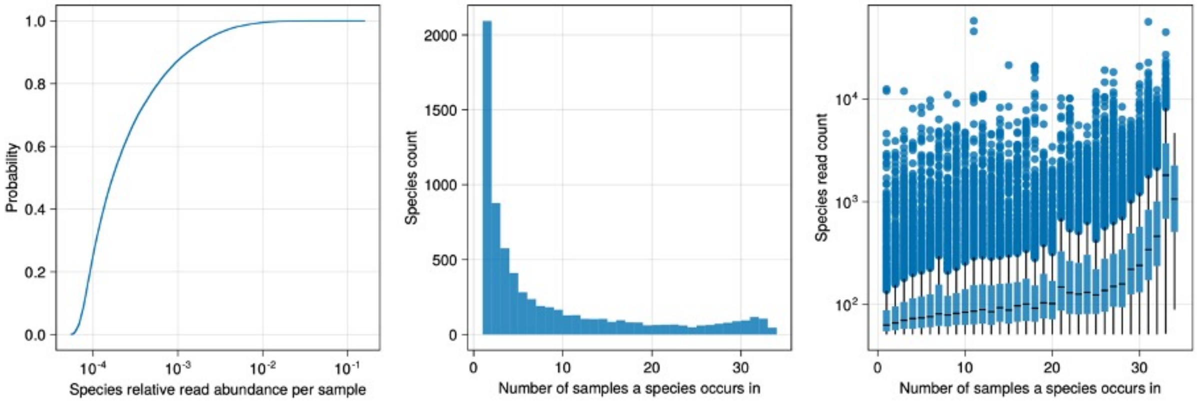


**Fig. S2.**
Species abundance distribution. From left to right: histogram of species read count for each occurrence in samples, histogram of number of occurrences in samples (1 to 34) for each species, and boxplot of species read count grouped by the number of occurrences in samples.

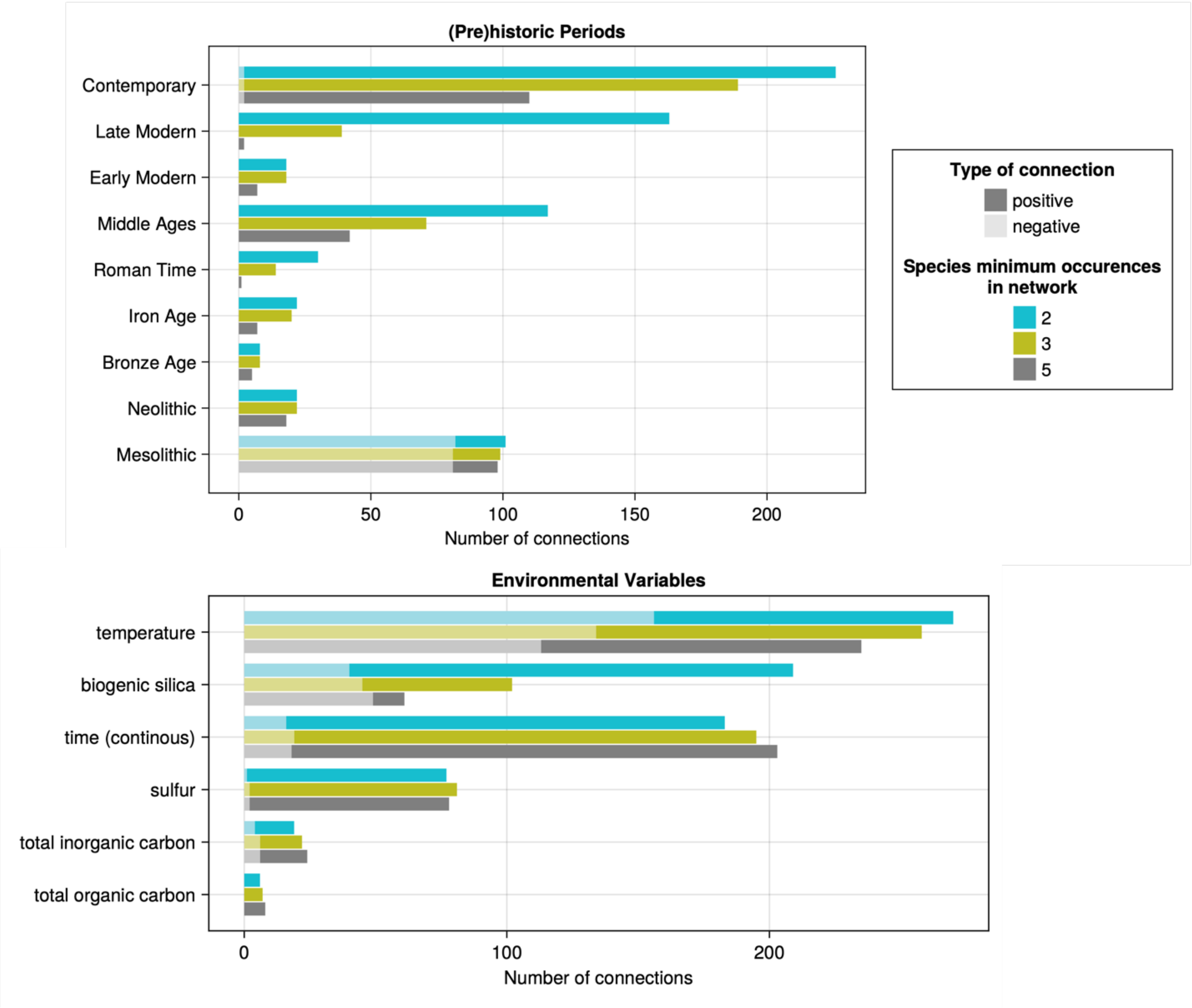


**Fig. S3.**

Number of connections, representing explanatory power of (pre)historic periods and selected environmental variables in networks, are plotted to show their association with the occurrences of microbes. Positive connections represent the increase of taxa. Univariate networks are analysed, so that both direct and indirect associations are considered. When networks were built with longer-term taxa and the pattern still holds, it implies that the effect of these time periods was long-lasting.

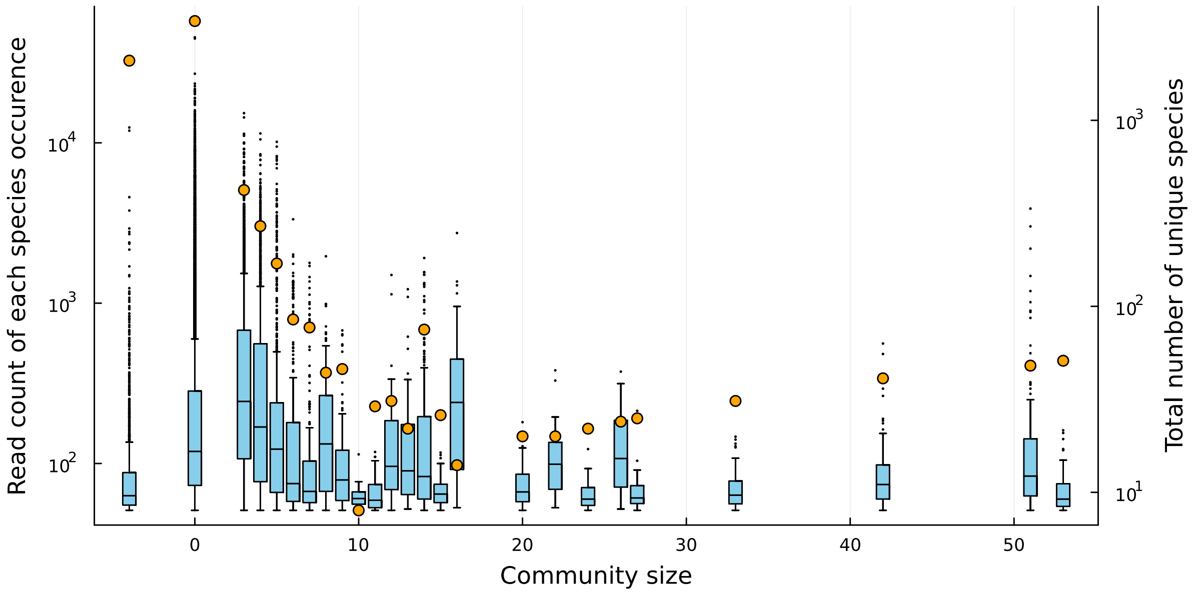


**Fig. S4.**
Characteristics of communities in the co-abundance network. Singletons are species that occur only in one sample, therefore not included in the network. Species with community size 0 are those in the network but are not members of any communities (size ≥3).

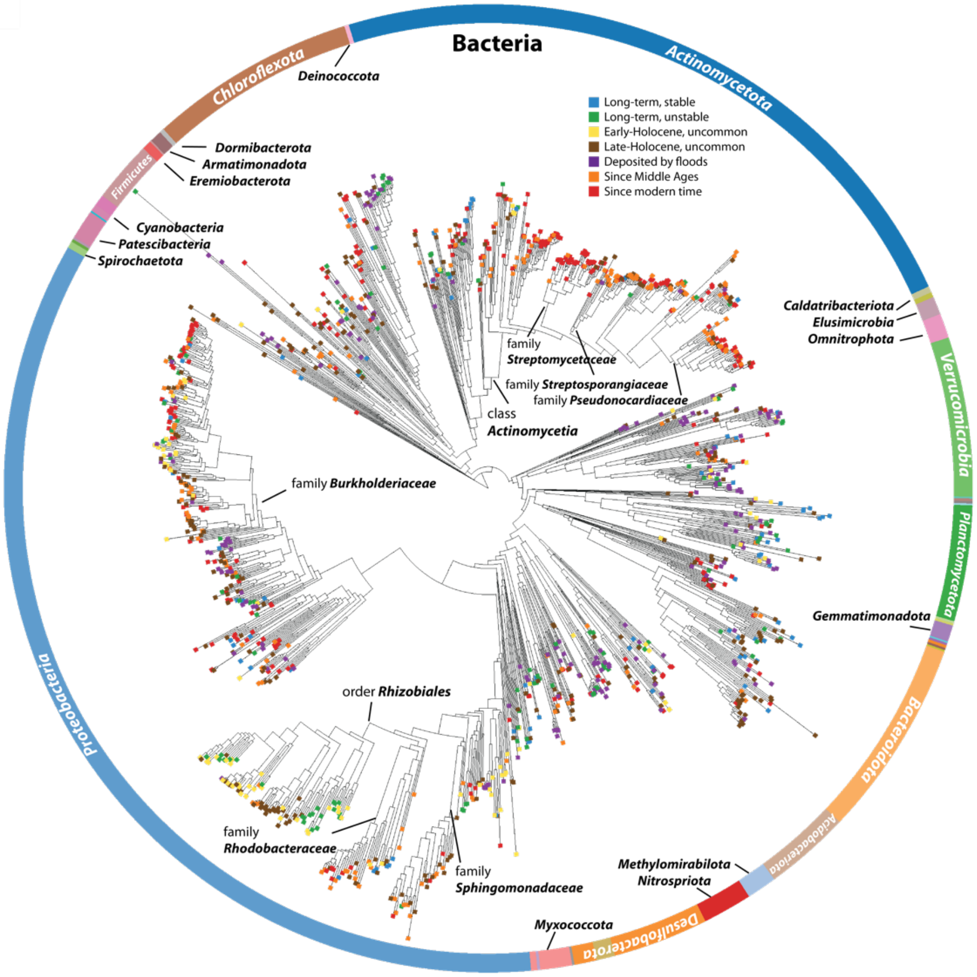


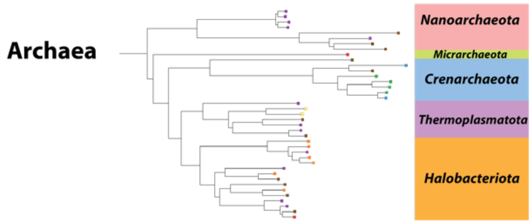


**Fig. S5.**
Taxonomic tree that includes bacterial and archaeal taxa in groups “Early-Holocene, uncommon” and “Late-Holocene, uncommon”.

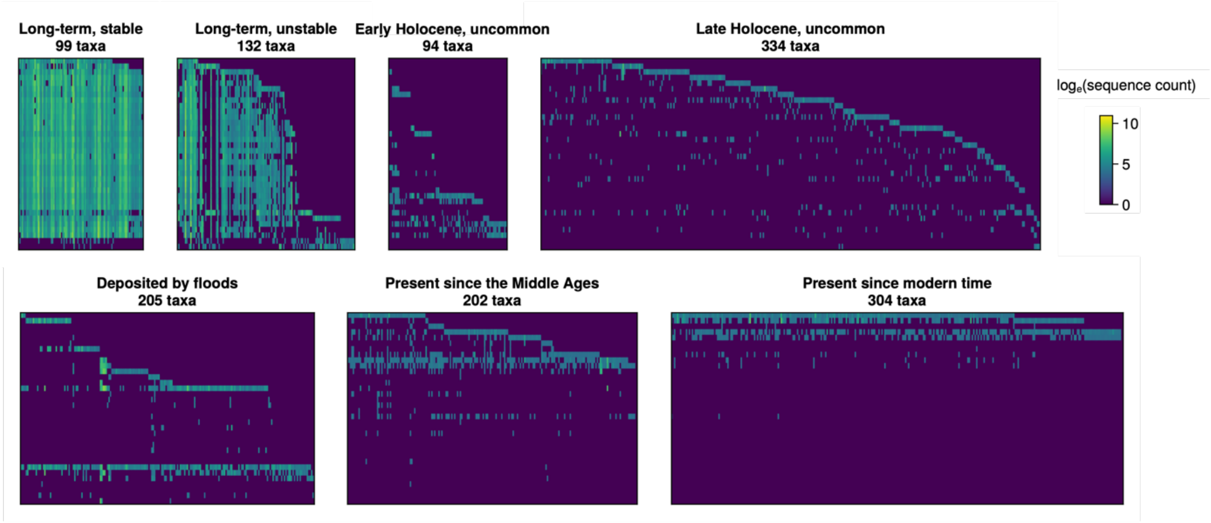


**Fig. S6.**
Heatmaps of sequence abundance for seven assemblage groups.

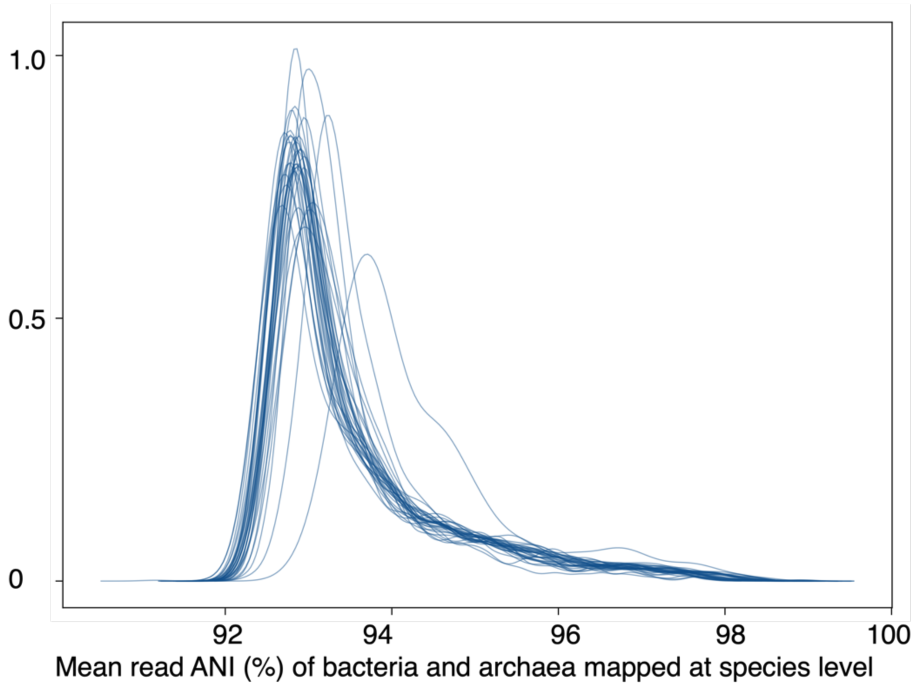


**Fig. S7.**
ANI distribution of samples. The outlier distribution is from HB_22, likely due to shallow sequencing depth.

**Table S1.**
Metadata of pooled libraries containing *sed*aDNA samples and controls.

| Library name | Library number | Sample name | Extraction number | Depth (cm) | Index PCR cycle | Molarity 130-330bp (pM) | Pooled volume (0.2 pmol) | p7 | p5 |
|---|---|---|---|---|---|---|---|---|---|
| AVXF-1-1 | YW03L.01 | HIBO_aDNA_118 | YW05E.02 | 102 | 4+4 | 74,625.60 | 2.68 | GTTGCGT | CCTGCGA |
| AVXF-1-2 | YW03L.02 | HIBO_aDNA_129 | YW05E.03 | 229 | 4+4 | 93,828.10 | 2.13 | TTGCGAA | TGCAGAG |
| AVXF-1-3 | YW03L.03 | HIBO_aDNA_7 | YW05E.04 | 408 | 4+4 | 86,359.60 | 2.32 | GTCGCAG | ACCTAGG |
| AVXF-1-4 | YW03L.04 | HIBO_aDNA_18 | YW05E.05 | 624.3 | 4+4 | 82,630.60 | 2.42 | CTAAGTA | TTGATCC |
| AVXF-1-5 | YW03L.05 | HIBO_aDNA_21 | YW05E.06 | 683.4 | 4+4 | 47,370.40 | 4.22 | AAGCTGA | ATCTTGC |
| AVXF-1-6 | YW03L.06 | HIBO_aDNA_29 | YW05E.07 | 845.6 | 4+4 | 48,315.00 | 4.14 | GATGCTG | TCTCCAT |
| AVXF-1-7 | YW03L.09 | EB | YW05E.10 | NA | 12+4 | 1,019.40 | 1.00 | GATTAGC | CCGGTAC |
| AVXF-1-8 | YW03L.10 | Bodensee/5-19 1 | YW03E.01 | 300.5 | 5+4 | 37,721.00 | 5.30 | AGAAGTC | GCCATAG |
| AVXF-1-9 | YW03L.11 | Bodensee/5-19 2 | YW03E.02 | 500.5 | 5+4 | 65,865.60 | 3.04 | ATAGTAC | GCCGGAC |
| AVXF-1-10 | YW03L.12 | Bodensee/5-19 3 | YW03E.03 | 700.5 | 4+4 | 32,136.60 | 6.22 | GATCTCG | CGTAGGC |
| AVXF-1-11 | YW03L.13 | Bodensee/5-19 4 | YW03E.04 | 900.5 | 4+4 | 36,347.70 | 5.50 | GGCTGCG | CTATGCC |
| AVXF-1-12 | YW03L.14 | Bodensee/5-19 5 | YW03E.05 | 1100.5 | 6+4 | 36,948.30 | 5.41 | GATCCAA | GAGTTAA |
| AVXF-1-13 | YW03L.15 | EB | YW03E.11 | NA | 12+4 | 5,803.10 | 1.00 | GAGCATG | CCGCTGG |
| AVXF-1-14 | YW03L.16 | Blank | NA | NA | 12+4 | 6,410.00 | 1.00 | TTGCGAA | CCTGCGA |
| AVXF-1-15 | YW04L.01 | HIBO_aDNA_17 | YW11E.01 | 605.5 | 9 | 15,612.90 | 12.81 | GTCGCAG | TGCAGAG |
| AVXF-1-16 | YW04L.02 | HIBO_aDNA_19 | YW11E.02 | 644.3 | 10 | 32,092.00 | 6.23 | CTAAGTA | ACCTAGG |
| AVXF-1-17 | YW04L.03 | HIBO_aDNA_27 | YW11E.03 | 804.9 | 10 | 17,290.10 | 11.57 | AAGCTGA | TTGATCC |
| AVXF-1-18 | YW04L.05 | HIBO_aDNA_35 | YW11E.06 | 966.1 | 10 | 25,714.50 | 7.78 | CCGCTGG | TCTCCAT |
| AVXF-1-19 | YW04L.06 | HIBO_aDNA_37 | YW11E.07 | 1003.8 | 8 | 20000* | 10.00 | ATATACG | CGGAGTT |
| AVXF-1-20 | YW04L.07 | HIBO_aDNA_39 | YW11E.08 | 1043.8 | 9 | 26,173.20 | 7.64 | GATTAGC | CAGGTCG |
| AVXF-1-21 | YW04L.10 | HIBO_aDNA_98 | YW11E.11 | 2383.1 | 7 | 36,377.50 | 5.50 | GATCTCG | GCCGGAC |
| AVXF-1-22 | YW04L.11 | HIBO_aDNA_67 | YW04E.04 | 1633.5 | 12 | 162,405.90 | 1.23 | GGCTGCG | CGTAGGC |
| AVXF-1-23 | YW04L.13 | EB | YW04E.12 | NA | 15 | 5,564.70 | 1.00 | GAGCATG | GAGTTAA |
| AVXF-1-24 | YW04L.14 | Blank | NA | NA | 15 | 4,784.00 | 1.00 | GTTGCGT | CCGCTGG |
| AVXF-1-25 | YW05L.01 | HIBO_aDNA_107 | YW12E.02 | 11 | 8 | 123,451.90 | 1.62 | GTCGCAG | CCTGCGA |
| AVXF-1-26 | YW05L.02 | HIBO_aDNA_111 | YW12E.03 | 35 | 8 | 94,048.50 | 2.13 | CTAAGTA | TGCAGAG |
| AVXF-1-27 | YW05L.03 | HIBO_aDNA_114 | YW12E.04 | 65 | 8 | 81,126.80 | 2.47 | AAGCTGA | ACCTAGG |
| AVXF-1-28 | YW05L.04 | HIBO_aDNA_116 | YW12E.05 | 85 | 9 | 74,864.40 | 2.67 | GATGCTG | TTGATCC |
| AVXF-1-29 | YW05L.05 | HIBO_aDNA_120 | YW12E.06 | 122 | 9 | 52,393.60 | 3.82 | CCGCTGG | ATCTTGC |
| AVXF-1-30 | YW05L.06 | HIBO_aDNA_123 | YW12E.07 | 152 | 9 | 60000* | 3.33 | ATATACG | TCTCCAT |
| AVXF-1-31 | YW05L.07 | HIBO_aDNA_126 | YW12E.08 | 182 | 9 | 54,037.50 | 3.70 | GATTAGC | CGGAGTT |
| AVXF-1-32 | YW05L.08 | HIBO_aDNA_128 | YW12E.09 | 209 | 9 | 47,563.80 | 4.20 | AGAAGTC | CAGGTCG |
| AVXF-1-33 | YW05L.09 | HIBO_aDNA_131 | YW12E.10 | 269 | 9 | 62,875.60 | 3.18 | ATAGTAC | CCGGTAC |
| AVXF-1-34 | YW05L.10 | HIBO_aDNA_9 | YW12E.11 | 447.5 | 9 | 81,988.00 | 2.44 | GATCTCG | GCCATAG |
| AVXF-1-35 | YW05L.11 | EB | YW12E.12 | NA | 11 | 106.4 | 1.00 | GGCTGCG | GCCGGAC |
| AVXF-1-36 | YW05L.12 | HIBO_aDNA_30 | YW11E.04 | 865.6 | 10 | 28,429.20 | 7.04 | GATCCAA | CGTAGGC |
| AVXF-1-37 | YW05L.13 | EB | YW11E.12 | NA | 11 | 2,777.00 | 1.00 | GAGCATG | CTATGCC |
| AVXF-1-38 | YW05L.15 | Bodensee/5-19 13 | YW02E.03 | 350.5 | 8 | 129,124.70 | 1.55 | TTGCGAA | CCGCTGG |
| AVXF-1-39 | YW05L.16 | Bodensee/5-19 14 | YW02E.04 | 550.5 | 8 | 125,460.70 | 1.59 | CTAAGTA | CCTGCGA |
| AVXF-1-40 | YW05L.17 | Bodensee/5-19 15 | YW02E.05 | 750.5 | 10 | 37,305.20 | 5.36 | AAGCTGA | TGCAGAG |
| AVXF-1-41 | YW05L.19 | EB | YW02E.12 | NA | 11 | 2,939.90 | 1.00 | CCGCTGG | TTGATCC |

| | | | | | | | | | |
|---|---|---|---|---|---|---|---|---|---|
| AVXF-1-42 | YW05L.20 | Blank | NA | NA | 11 | NA | 1.00 | ATATACG | ATCTTGC |
| AVXF-1-43 | YW06L.01 | HIBO_aDNA_100 | YW12E.01 | 0 | 8 | 146,904.50 | 1.36 | GATTAGC | TCTCCAT |
| AVXF-1-44 | YW06L.02 | Blank | NA | NA | 11 | 6,857.60 | 1.00 | AGAAGTC | CGGAGTT |

* Calibrated value based on Bioanalyzer estimation.

**Table S2.**
Illumina sequencing data report.

| Library ID | Library type | Yield (Mb) | %PF | Cluster (PF) | Q30 | Mean qual. (PF) |
|---|---|---|---|---|---|---|
| AVXF-1-2 | sample | 15437 | 100 | 77184410 | 92.2 | 35.53 |
| AVXF-1-11 | sample | 14185 | 100 | 70925328 | 94.79 | 36.1 |
| AVXF-1-5 | sample | 14057 | 100 | 70282616 | 93.09 | 35.69 |
| AVXF-1-40 | sample | 14055 | 100 | 70274719 | 92.37 | 35.53 |
| AVXF-1-6 | sample | 13778 | 100 | 68889610 | 92.8 | 35.64 |
| AVXF-1-36 | sample | 13730 | 100 | 68649711 | 92.18 | 35.46 |
| AVXF-1-15 | sample | 13477 | 100 | 67384276 | 91.93 | 35.42 |
| AVXF-1-28 | sample | 13454 | 100 | 67269837 | 92.32 | 35.53 |
| AVXF-1-30 | sample | 13303 | 100 | 66513960 | 92.07 | 35.45 |
| AVXF-1-27 | sample | 13122 | 100 | 65608610 | 93.53 | 35.79 |
| AVXF-1-8 | sample | 12937 | 100 | 64685965 | 92.7 | 35.59 |
| AVXF-1-31 | sample | 12827 | 100 | 64135222 | 92.66 | 35.59 |
| AVXF-1-32 | sample | 12781 | 100 | 63906376 | 92.3 | 35.5 |
| AVXF-1-33 | sample | 12781 | 100 | 63904738 | 92.03 | 35.44 |
| AVXF-1-25 | sample | 12391 | 100 | 61956963 | 91.87 | 35.38 |
| AVXF-1-12 | sample | 12358 | 100 | 61788565 | 92.26 | 35.5 |
| AVXF-1-10 | sample | 12114 | 100 | 60570833 | 92.7 | 35.58 |
| AVXF-1-43 | sample | 11909 | 100 | 59544209 | 91.9 | 35.41 |
| AVXF-1-3 | sample | 11849 | 100 | 59244258 | 93.91 | 35.87 |
| AVXF-1-1 | sample | 11801 | 100 | 59007284 | 92.46 | 35.53 |
| AVXF-1-9 | sample | 11716 | 100 | 58577891 | 93.19 | 35.69 |
| AVXF-1-39 | sample | 11552 | 100 | 57762124 | 92.86 | 35.63 |
| AVXF-1-21 | sample | 11119 | 100 | 55593145 | 94.9 | 36.12 |
| AVXF-1-4 | sample | 10883 | 100 | 54414602 | 94.46 | 36 |
| AVXF-1-20 | sample | 10673 | 100 | 53366366 | 93.49 | 35.78 |
| AVXF-1-19 | sample | 10512 | 100 | 52559211 | 92.73 | 35.6 |
| AVXF-1-17 | sample | 10325 | 100 | 51627231 | 91.69 | 35.36 |
| AVXF-1-16 | sample | 10031 | 100 | 50154558 | 91.92 | 35.44 |
| AVXF-1-38 | sample | 9760 | 100 | 48799055 | 93.43 | 35.81 |
| AVXF-1-29 | sample | 9685 | 100 | 48425607 | 90.08 | 35.06 |
| AVXF-1-34 | sample | 9098 | 100 | 45488093 | 92.81 | 35.61 |
| AVXF-1-18 | sample | 8944 | 100 | 44717514 | 90.59 | 35.16 |
| AVXF-1-26 | sample | 8905 | 100 | 44524257 | 93.82 | 35.86 |
| AVXF-1-22 | sample | 1776 | 100 | 8881398 | 94.45 | 36.01 |
| AVXF-1-14 | control | 96 | 100 | 481385 | 73.48 | 31.37 |
| AVXF-1-13 | control | 89 | 100 | 446487 | 83.5 | 33.56 |
| AVXF-1-7 | control | 65 | 100 | 323536 | 84.15 | 33.7 |
| AVXF-1-24 | control | 56 | 100 | 280586 | 85.77 | 34.16 |
| AVXF-1-23 | control | 40 | 100 | 201373 | 87.88 | 34.57 |
| AVXF-1-44 | control | 37 | 100 | 186932 | 70.11 | 30.52 |
| AVXF-1-41 | control | 9 | 100 | 43394 | 87.08 | 34.48 |
| AVXF-1-42 | control | 7 | 100 | 35067 | 86.21 | 34.26 |

| | | | | | | |
|---|---|---|---|---|---|---|
| AVXF-1-35 | control | 5 | 100 | 25198 | 87.33 | 34.54 |
| AVXF-1-37 | control | 1 | 100 | 5529 | 81.04 | 32.97 |

**Table S3.**
Radiocarbon dates of HIBO19 sediment core measured in this study.

| sample name | sample label | composite depth top(cm) | composite depth bottom(cm) | depth in model (cm) | included in model | age (y) | +- (y) | weight (ugC) | sample collector |
|---|---|---|---|---|---|---|---|---|---|
| Hipercorig_14C_13 | HB14C_13 | 42.2 | 43.2 | 42.7 | TRUE | 150 | 70 | 94 | Wang |
| Hipercorig_14C_01 | HB14C_01 | 110 | 111 | 110.5 | TRUE | 458 | 69 | 83 | Wang |
| Hipercorig_14C_20 | HB14C_20 | 198.5 | 199.5 | 199 | TRUE | 952 | 79 | 45 | Wang |
| Hipercorig_14C_03 | HB14C_03 | 273 | 274 | 273.5 | TRUE | 1,764 | 74 | 62 | Wang |
| Hipercorig_14C_14 | HB14C_14 | 301.5 | 302.5 | 302 | TRUE | 2,082 | 96 | 146 | Wang |
| Hipercorig_14C_15 | HB14C_15 | 363 | 364 | 363.5 | TRUE | 2,703 | 94 | 84 | Wang |
| Hipercorig_14C_16 | HB14C_16 | 423.5 | 424.5 | 424 | TRUE | 3,191 | 151 | 20 | Wang |
| Hipercorig_14C_05 | HB14C_05 | 477.7 | 478.7 | 478.2 | TRUE | 3,665 | 85 | 32 | Wang |
| Hipercorig_14C_19 | HB14C_19 | 531.7 | 532.7 | NA | FALSE | 5,963 | 204 | 15 | Wang |
| Hipercorig_14C_08 | HB14C_08 | 761.9 | 762.9 | 762.4 | TRUE | 6,798 | 100 | 65 | Wang |
| Hipercorig_14C_17 | HB14C_17 | 820.9 | 821.9 | 821.4 | TRUE | 7,861 | 237 | 26 | Wang |
| Hipercorig_14C_09 | HB14C_09 | 1202.7 | 1203.7 | NA | FALSE | 35,218 | 743 | 94 | Wang |
| Hipercorig_14C_10 | HB14C_10 | 1420.6 | 1421.6 | NA | FALSE | 12,637 | 128 | 98 | Wang |
| Hipercorig_14C_11 | HB14C_11 | 1722.6 | 1723.6 | NA | FALSE | 13,290 | 158 | 60 | Wang |
| Hipercorig_14C_12 | HB14C_12 | 1996.7 | 1997.7 | NA | FALSE | 12,794 | 146 | 99 | Wang |
| 14C-1a | 14C-1a | 883 | 885 | 884 | TRUE | 8322 | 92 | 164 | Schaller |
| 14C-2a | 14C-2a | 902 | 912 | 911 | TRUE | 8434 | 191 | 23 | Schaller |
| 14C-3a | 14C-3a | 963 | 965 | 964 | TRUE | 9598 | 41 | 514 | Schaller |
| 14C-7 | 14C-7 | 2397.5 | 2398.5 | 1328 | TRUE | 11772 | 72 | 188 | Schaller |

**Table S4.**
A summary table of sequencing read counts per library after each filtering step.

| Library name | Library short name | Number of raw reads | Number of paired reads after fastp | Read count after sga low-complex dust=4 | Read count after sga remove dupplicates | Reads after prinseq++ trim both ends ployA/T, derep and minLen30 |
|---|---|---|---|---|---|---|
| AVXF-1-01 | HB_01 | 118,014,568 | 46,039,695 | 45,216,476 | 35,591,373 | 34,439,141 |
| AVXF-1-02 | HB_02 | 154,368,820 | 61,248,284 | 60,508,355 | 48,163,970 | 42,872,398 |
| AVXF-1-03 | HB_03 | 118,488,516 | 49,152,003 | 48,599,298 | 38,633,494 | 36,836,484 |
| AVXF-1-04 | HB_04 | 108,829,204 | 43,523,085 | 43,176,658 | 34,256,884 | 33,315,999 |
| AVXF-1-05 | HB_05 | 140,565,232 | 54,229,512 | 53,706,441 | 42,588,703 | 39,553,771 |
| AVXF-1-06 | HB_06 | 137,779,220 | 54,121,536 | 53,673,393 | 39,713,558 | 37,591,024 |
| AVXF-1-07 | HB_07 | 647,072 | 133,773 | 132,639 | 99,924 | 99,535 |
| AVXF-1-08 | HB_08 | 129,371,930 | 51,624,698 | 50,664,286 | 39,891,324 | 37,756,500 |
| AVXF-1-09 | HB_09 | 117,155,782 | 48,683,531 | 48,013,661 | 38,271,263 | 36,587,054 |
| AVXF-1-10 | HB_10 | 121,141,666 | 47,977,414 | 47,437,032 | 37,930,996 | 36,302,481 |
| AVXF-1-11 | HB_11 | 141,850,656 | 53,582,423 | 53,434,651 | 42,586,142 | 39,378,841 |
| AVXF-1-12 | HB_12 | 123,577,130 | 47,841,689 | 47,215,668 | 37,570,970 | 36,080,679 |
| AVXF-1-13 | HB_13 | 892,974 | 127,734 | 127,137 | 92,762 | 92,446 |
| AVXF-1-14 | HB_14 | 962,770 | 81,614 | 81,042 | 67,505 | 67,189 |
| AVXF-1-15 | HB_15 | 134,768,552 | 51,596,267 | 50,588,457 | 39,805,655 | 37,735,366 |
| AVXF-1-16 | HB_16 | 100,309,116 | 38,577,547 | 37,809,096 | 29,965,282 | 29,545,431 |
| AVXF-1-17 | HB_17 | 103,254,462 | 38,712,923 | 38,134,010 | 29,113,786 | 28,771,517 |
| AVXF-1-18 | HB_18 | 89,435,028 | 33,143,597 | 32,598,368 | 24,920,086 | 24,754,693 |
| AVXF-1-19 | HB_19 | 105,118,422 | 40,443,263 | 40,042,125 | 31,688,176 | 31,151,526 |
| AVXF-1-20 | HB_20 | 106,732,732 | 43,019,187 | 42,580,574 | 34,783,945 | 33,862,796 |
| AVXF-1-21 | HB_21 | 111,186,290 | 39,177,701 | 39,106,471 | 27,269,436 | 26,610,929 |
| AVXF-1-22 | HB_22 | 17,762,796 | 5,380,318 | 5,356,889 | 4,448,870 | 4,407,198 |
| AVXF-1-23 | HB_23 | 402,746 | 68,029 | 67,816 | 54,105 | 53,967 |
| AVXF-1-24 | HB_24 | 561,172 | 12,431 | 12,325 | 10,849 | 10,800 |
| AVXF-1-25 | HB_25 | 123,913,926 | 46,332,035 | 45,926,604 | 33,594,099 | 32,797,003 |
| AVXF-1-26 | HB_26 | 89,048,514 | 35,549,980 | 35,078,871 | 28,618,610 | 28,322,023 |
| AVXF-1-27 | HB_27 | 131,217,220 | 53,792,228 | 53,112,207 | 42,173,905 | 39,368,322 |
| AVXF-1-28 | HB_28 | 134,539,674 | 54,126,817 | 52,882,995 | 41,062,597 | 38,620,990 |
| AVXF-1-29 | HB_29 | 96,851,214 | 36,225,797 | 35,240,358 | 28,175,771 | 27,871,769 |
| AVXF-1-30 | HB_30 | 133,027,920 | 52,185,624 | 50,773,926 | 39,702,509 | 37,642,117 |
| AVXF-1-31 | HB_31 | 128,270,444 | 51,402,057 | 50,525,094 | 41,034,175 | 38,603,771 |
| AVXF-1-32 | HB_32 | 127,812,752 | 50,625,112 | 49,579,601 | 40,658,116 | 38,345,732 |
| AVXF-1-33 | HB_33 | 127,809,476 | 49,695,847 | 48,520,888 | 35,905,159 | 34,751,561 |
| AVXF-1-34 | HB_34 | 90,976,186 | 36,736,089 | 36,087,784 | 29,106,515 | 28,764,893 |
| AVXF-1-35 | HB_35 | 50,396 | 15,327 | 15,247 | 14,686 | 14,515 |
| AVXF-1-36 | HB_36 | 137,299,422 | 52,770,322 | 51,959,824 | 39,675,013 | 37,640,420 |
| AVXF-1-37 | HB_37 | 11,058 | 1,957 | 1,935 | 1,702 | 1,695 |
| AVXF-1-38 | HB_38 | 97,598,110 | 39,358,810 | 39,053,911 | 31,088,918 | 30,546,584 |
| AVXF-1-39 | HB_39 | 115,524,248 | 48,160,553 | 47,619,507 | 39,115,499 | 37,241,515 |
| AVXF-1-40 | HB_40 | 140,549,438 | 53,960,454 | 53,204,142 | 40,299,830 | 37,938,011 |
| AVXF-1-41 | HB_41 | 86,788 | 30,823 | 30,305 | 28,979 | 28,869 |

| | | | | | | |
|---|---|---|---|---|---|---|
| AVXF-1-42 | HB_42 | 70,134 | 19,344 | 19,013 | 18,323 | 18,235 |
| AVXF-1-43 | HB_43 | 119,088,418 | 44,158,613 | 43,931,591 | 35,184,837 | 34,172,063 |
| AVXF-1-44 | HB_44 | 373,864 | 26,109 | 25,694 | 24,845 | 24,743 |

**Supplementary File**

Fragment size distribution of pooled *sed*aDNA library and individual sample/control libraries. Lengths shown in the figure include adapters (~130 bp in length), therefore the lengths of original DNA reads peak at ~169-130=39 bp.

**Data S1. (separate file)**
Data of assemblage group "Long-term, stable". **a.** Table of DNA sequence counts. **b.** Table of DNA damage. **c.** Taxonomic information of identified taxa.

**Data S2. (separate file)**
Data of assemblage group "Long-term, unstable". **a.** Table of DNA sequence counts. **b.** Table of DNA damage. **c.** Taxonomic information of identified taxa.

**Data S3. (separate file)**
Data of assemblage group "Early-Holocene, uncommon". **a.** Table of DNA sequence counts. **b.** Table of DNA damage. **c.** Taxonomic information of identified taxa.

**Data S4. (separate file)**
Data of assemblage group "Late-Holocene, uncommon". **a**. Table of DNA sequence counts. **b.** Table of DNA damage. **c.** Taxonomic information of identified taxa.

**Data S5. (separate file)**
Data of assemblage group "Deposited by floods". **a**. Table of DNA sequence counts. **b.** Table of DNA damage. **c.** Taxonomic information of identified taxa**.**

**Data S6. (separate file)**
Data of assemblage group "Present since the Middle Ages". **a.** Table of DNA sequence counts. **b.** Table of DNA damage. **c.** Taxonomic information of identified taxa.

**Data S7. (separate file)**
Data of assemblage group "Present since modern time". **a.** Table of DNA sequence counts. **b.** Table of DNA damage. **c.** Taxonomic information of identified taxa.
